# Single-cell visual proteomics of a minimal bacterium reveals structural coordination in gene expression

**DOI:** 10.1101/2025.10.13.682074

**Authors:** Joseph M. Dobbs, Rasmus K. Jensen, Julia Mahamid

**Affiliations:** Molecular Systems Biology Unit, European Molecular Biology Laboratory (EMBL), Heidelberg, Germany; Collaboration for joint PhD degree between EMBL and Heidelberg University, Faculty of Biosciences, Heidelberg, Germany; Cell Biology and Biophysics Unit, EMBL, Heidelberg, Germany

## Abstract

Translation is a central process in gene expression. Regulation of this process is complex, depends on factors that include cell state and the subcellular environment, and is subject to modulation via crosstalk to processes like transcription or translocation. Here, we used cryo-electron tomography of native and antibiotic-perturbed *Mycoplasma pneumoniae* cells to resolve 140 maps that recapitulate bacterial translation during the initiation, elongation, termination, and recycling phases. We visualized multiple transcription-translation complexes, allowing us to propose a threading-based translation reinitiation mechanism, and to provide structural evidence for a long-hypothesized supercomplex that coordinates transcription, translation, and membrane attachment. We further resolved abundant membrane-associated large ribosomal subunits, and suggest that dissociation from membranes depends on the conditional initiation of new translation, consistent with a conserved mechanism in mammalian cells. This work visualizes the multilayered control of bacterial translation and demonstrates the power of in-cell structural biology to investigate regulatory circuits in gene expression.

## Introduction

Ribosomes are conserved molecular machines that translate messenger RNA (mRNA), transcribed by RNA polymerase (RNAP), into proteins. In bacteria, the 70S ribosome assembles from a 50S large subunit and a 30S small subunit. The translation process starts with recognition of the start codon on an mRNA by the 30S during the initiation phase, followed by joining of the 50S and elongation of the nascent peptide, termination at the stop codon, and recycling of the ribosomal subunits in preparation for a new round of translation. While this essential process has been studied at great depth in prokaryotes, many steps, and the interplay between layers of regulation across the four phases of translation, retain substantial uncertainty.^1,2^

This is particularly evident in the initiation phase, where the sequence of regulatory events is unclear. In vitro, initiation factors (IFs) 1,2, and 3, the initiating N-formylmethionine transfer RNA (fMet-tRNA^fMet^, or initiator tRNA) and the mRNA may bind in any order to the 30S.^3,4^ However, single-molecule fluorescence experiments with factors purified from *Escherichia coli*^3,5,6^ favor an order of assembly where IF2 and/or IF3 binding to the 30S occurs first, followed by IF1, and finally the initiator tRNA. Structural studies have further proposed a pathway involving mRNA binding before the tRNA,^7^ as recruitment of tRNA could physically block mRNA delivery to the initiation-competent 30S. This mRNA delivery is thought to rely on binding of the Shine-Dalgarno (SD) sequence in the mRNA 5’ untranslated region (5’ UTR) to the complimentary anti-Shine-Dalgarno (aSD) sequence on the 30S to accommodate the mRNA, with the associated start codon, into the complex.^8–10^

Translation in prokaryotes is further influenced by the lack of a nucleus, allowing mRNA and protein synthesis to take place in the same cellular compartment. An additional regulatory mechanism thus emerges that typically manifests itself in tight coordination between the elongation rates of transcription and translation.^11–14^ The direct association of ribosomes and RNAPs in transcription-translation complexes (T-TCs), also known as expressomes, has been characterized by structural studies of reconstituted *E. coli* complexes^13–19^ and in intact *Mycoplasma pneumoniae* cells during translation elongation.^20^ More recently, reconstitution studies have decoded interactions between the two machineries during translation initiation in *E. coli* and *Thermus thermophilus*,^21–23^ and highlight important roles for the SD sequence and ribosomal protein S1 during mRNA delivery. However, in many cases, mRNA delivery and accommodation in prokaryotic initiation does not strictly depend on classical SD-aSD interactions.^24–26^ The proportion of SD-containing open reading frames (ORFs) markedly varies across species, with some clades, including the complete phylum *Bacteroidetes*,^24–28^ lacking them entirely. In *M. pneumoniae*, only 8% of ORFs contain SD sequences (54% in *E. coli*),^26,29^ and the majority of these occur outside of the 5’ UTRs where they are canonically productive. Even in bacteria with abundant SD sequences, these are not required to properly position 30S complexes: disruption of SD/aSD pairing in an *E. coli* screen demonstrated that ribosomes can still select correct initiation sites.^30^ Likewise, while ribosomal protein S1 is important for mRNA recruitment,^22,31,32^ it is poorly conserved outside of Gram-negative bacteria^33^ and is absent in *M. pneumoniae*.^34^ In fact, *E. coli* ribosomes lacking S1 retain nearly wild-type levels of translation activity in vitro.^35^ Therefore, alternative mechanisms for translation initiation must exist, likely involving additional regulatory layers that are yet to be characterized.

After the start codon is recognized in the 30S, the 50S binds, IFs dissociate, and the assembled 70S ribosome elongates the nascent peptide in a process that requires peptidyl-tRNAs and elongation factors (EFs) G and Tu.^1,2,4^ Elongation terminates at the stop codon and dependent on one or more release factors (RFs), ribosome recycling factor (RRF), and EFG,^1,2,4^ the nascent protein is released and the subunits are recycled to restart the translation cycle.

While most cellular proteins are cytoplasmic, many are targeted to the cell membrane or secreted, and are thus subject to additional regulation. In prokaryotes, translocation across the membrane occurs either post-translationally, where fully translated proteins are moved through a membrane channel, or co-translationally, where recruitment of the ribosome to a membrane channel allows translocation during peptide synthesis.^36^ In co-translational translocation, targeting to the membrane is accomplished by recognition of the nascent peptide’s signal sequence by the signal recognition particle (SRP), and ribosomes can then directly interact with the translocation machinery including SecDF and the SecYEG channel.^37–41^ Importantly, in cells, this pathway takes place within the context of additional regulatory dimensions: processes like transcription-translation coupling could occur simultaneously, and it is unclear how the different phases of translation, and particularly the recycling of components, are regulated at the membrane.

A number of recent in-cell structural studies have begun to provide a comprehensive view of the translation elongation cycle in diverse species,^42–51^ but have yet to develop a complete cellular view of translation across its different phases. This stems from technical challenges in reliable and complete localization of complexes smaller than fully assembled 70S ribosomes within the crowded intracellular environment, and in the complexity of subsequent structural analyses given the expected large variability in translation complexes. Here, we comprehensively mapped and analyzed 30S, 50S, and 70S ribosomes in individual cells of the minimal bacterium *M. pneumoniae* imaged with cryogenic electron tomography (cryo-ET). We quantified complexes across the four translation phases and visualized their coordination with other cellular processes. By introducing roadblocks in transcription or translation with targeted antibiotic perturbations, stabilizing rare and transient complexes, we further dissected the molecular wiring of translational control at different regulatory levels and at single-cell resolution.

## Results and Discussion

### The coordination of translation complexes with cell volume

Our previous work characterized elongating 70S ribosomes in detail and at high resolution in *M. pneumoniae* cells.^20,45^ Here, we aimed for a comprehensive analysis of all translational complexes, including the large and small ribosomal subunits, to realize an understanding of the translation system as a whole in single cells during normal cellular growth. We therefore employed an exhaustive localization scheme and computational classification of complexes in tomograms of 254 native cells (Fig. 1A, B, Fig. S1A, B Methods), recovering a total of 29,654 free 30S subunits (Fig. 1C), 27,655 free 50S subunits (Fig. 1D) and 51,496 assembled 70S ribosomes (Fig. 1E). We mapped the complexes back into their intracellular positions on a per-tomogram basis (Fig. 1A, B), and combined those with quantification of cell volumes to derive their concentrations (Fig. 1F, Fig. S1C, Methods).

**Figure 1:**
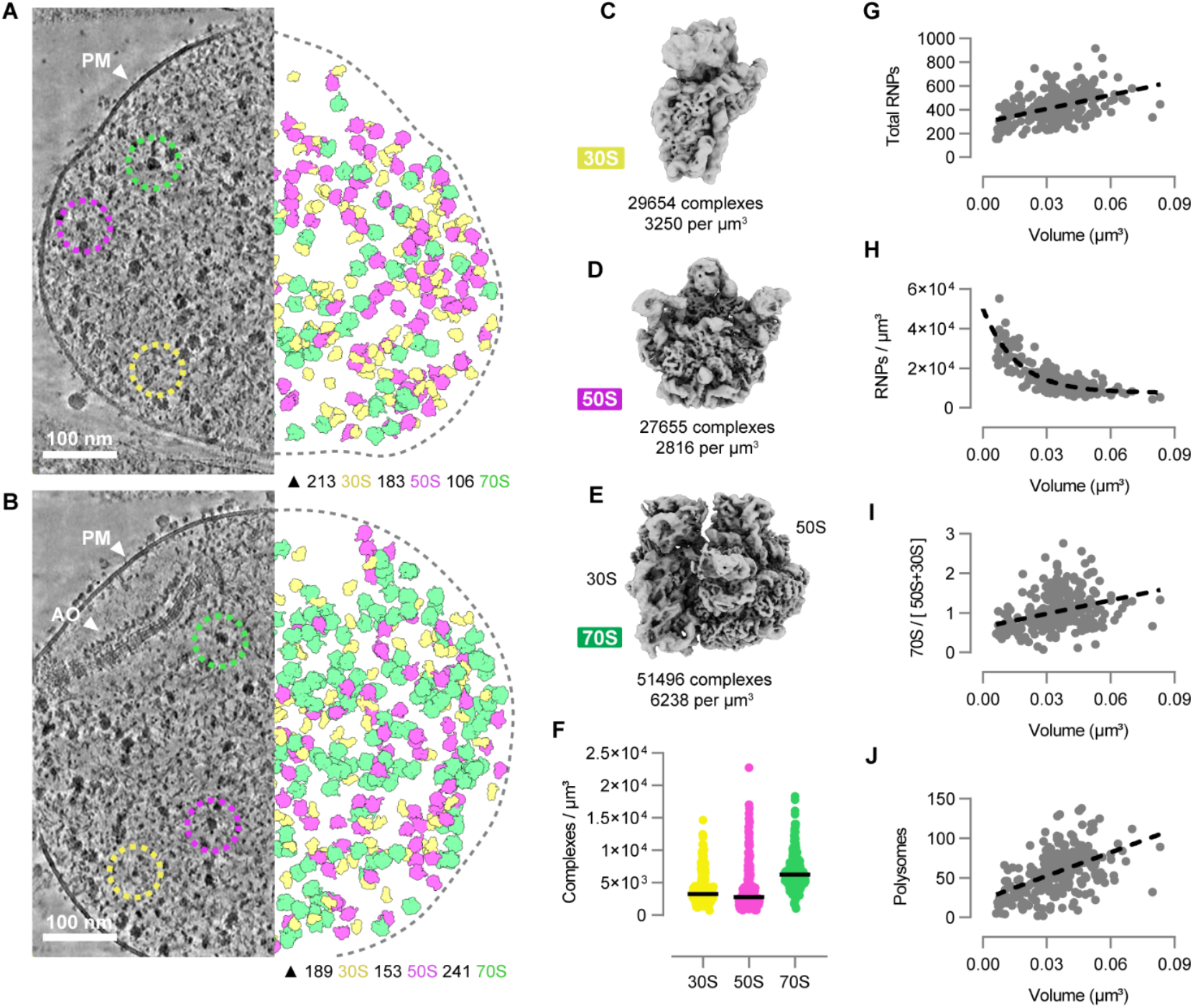
Ribosomal complex counts and assembly state correlate with cell volume. **(A, B)** Denoised tomographic slices of native *M. pneu-moniae* cells (left) and corresponding annotations (right) of 70S ribosomes (green), 50S large subunits (magenta) and 30S small subunits (yellow). Cell in **(A)** exhibited a lower proportion of assembled ribosomes than the cell in **(B).** PM: plasma membrane. AO: attachment organelle. **(C-E)** Consensus maps of the recovered **(C)** 30S (refined to 4.8 Å), **(D)** 50S (4.2 Å), and **(E)** 70S (3.8 Å). **(F-J)** Per-cell RNP quantifications related to cell volume. **(F)** Quantification of RNPs per µm^3^. Lines represent mean. **(G)** Total number of RNPs as a function of volume, fitted with a linear curve (R^2^ 0.1859). **(H)** RNP concentration per µm^3^ as a function of volume, fitted with an exponential decay curve (RMSE 4x10^3^). **(I)** Ratio of assembled 70S ribosomes to free 50S and 30S subunits as a function of volume, fitted with a linear curve (R^2^ 0.1069). **(J)** Plot of polysome count (distance criterion 7 nm) as a function of volume, fitted with a linear curve (R^2^ 0.2456). Dots in plots represent individual native cells (n=254). Related to Figure S1.

We noted extensive heterogeneity in the distributions of fully assembled ribosomes and free subunits across the cell population (Fig. 1A, B, F). We hypothesized that this heterogeneity may be related to cell cycle stage, which cannot be synchronized in *M. pneumoniae*, and therefore investigated the relationship between cell volume (as a proxy for growth) and per-cell counts of the 30S, 50S and 70S complexes, collectively referred to here as ribonucleoprotein particles (RNPs). This analysis showed that as cell volume increased, the sum total of RNPs in each cell increased linearly (Fig. 1G), but RNP concentration decreased exponentially (Fig. 1H). As a second proxy for cell cycle stage, we examined the number of attachment organelles, which replicate concurrently with genome duplication,^52^ and found that RNP counts were higher in cells with >1 attachment organelle (Fig. S1D). Thus, RNP production increases during cell growth, but does not keep pace with the physical expansion of the cell. Furthermore, we observed a linear relationship between cell volume and the ratio of assembled ribosomes to free subunits (Fig. 1I), and a linear increase in polysome counts with cell volume (Fig. 1J, Fig. S1E, Methods), suggesting a greater proportion of active translation in larger cells. In agreement with these results, a linear relationship between ribosome count and cell size, and the requirement for increased protein synthesis during cellular growth,^53–58^ are well established. Our results thus illustrate a strong correlation between translation regulation and cell state.

### Visualizing and quantifying complexes along the complete translation cycle

To disentangle the complexity of translation regulation within the cell population, we next performed extensive structural subclassification of the RNPs (Fig. S2, Methods). This analysis recapitulated the canonical translation cycle, recovering eight complexes in the initiation phase, fourteen in elongation, one in termination, and three in recycling, each quantified by their median abundance per cell (Fig. 2A, total fractions provided in Table S1). Additional complexes tangential to this cycle, including those found in crosstalk to other cellular machineries, were also resolved and quantified (Table S1). Here, we first describe our overall findings, and in later sections expand on critical stages of translational control.

**Figure 2:**
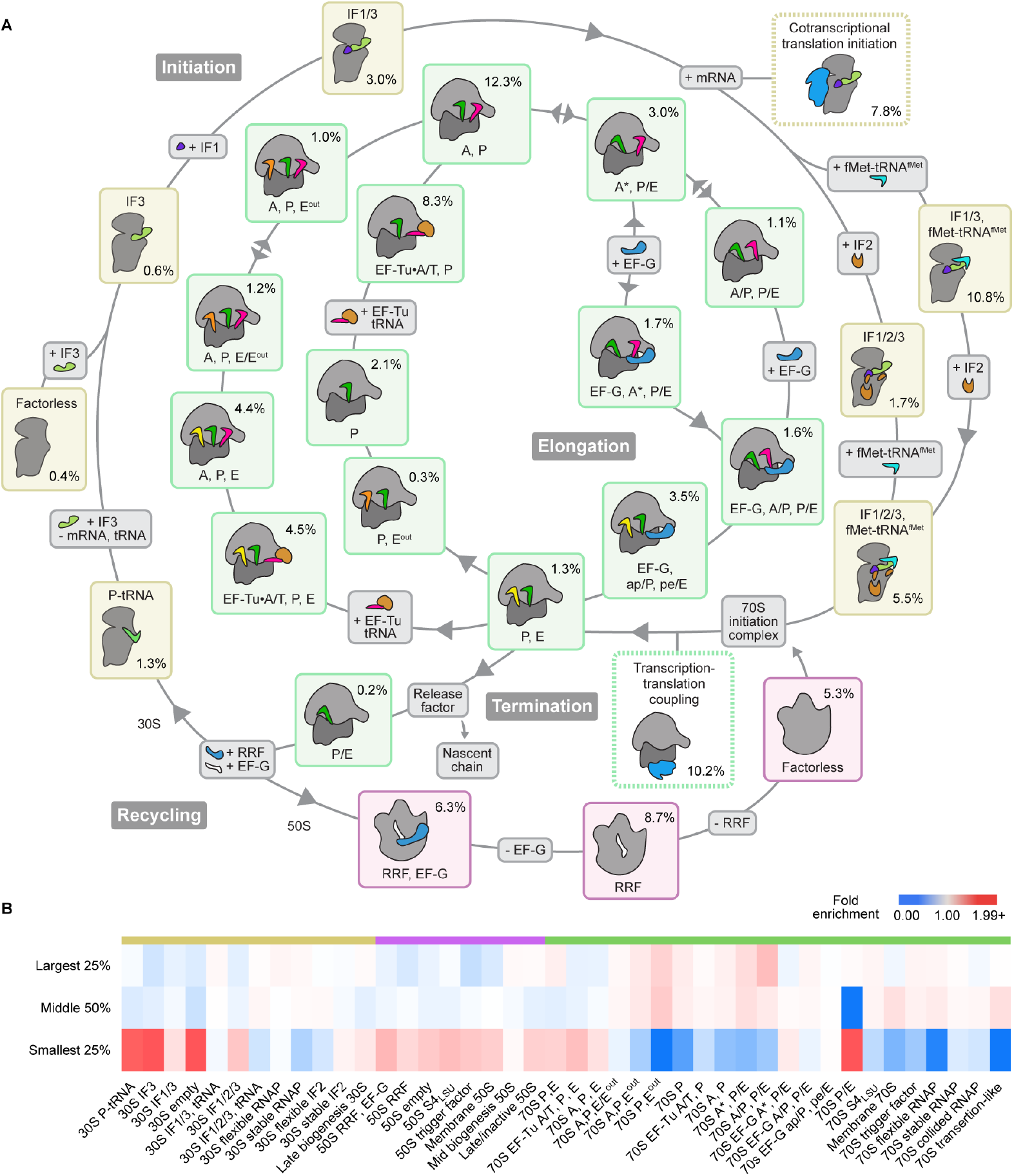
Classification recapitulates and quantifies the translational control cycle. **(A)** Schematic of the translation cycle in native cells. Each cartoon represents a complex resolved by classification from *M. pneumoniae*, and the median per-cell percentages of all RNPs are listed. Table S1 lists total fractions. **(B)** Abundances of classified translation complexes represented by median fold enrichment over abundance in the total population, shown separately for the largest 25% of cells by volume (n=64), the middle 50% (n=126), or the smallest 25% (n=64). Coloured bars (top) relate to the consensus refinement from which the class is derived (30S, 50S or 70S), as shown in (A). Related to Figures S2 and S3, and Table S1.

Resolving for the first time bacterial translation initiation directly in cells, we quantified 30S subunits and their association with initiation factors and initiator tRNAs (Fig. 2A: yellow boxes, Table S1). We describe the derived sequence of initiation in the following section. We did not resolve inactive 30S complexes, characterized by helix 44 repositioning, which have been suggested to account for the majority of small subunits in *E. coli*.^59,60,22^ Instead, we found that at least 89% of 30S complexes were bound by initiation factors.

Translation elongation in *M. pneumoniae* was analyzed in depth in our previous work.^45^ Here, we used an expanded classification strategy (Methods) to uncover multiple elongation states previously unresolved in the native bacterium, including 70S ribosomes (Fig. 2A: green boxes) with simultaneous engagement of A-, P- and E-site tRNAs (4.4% of RNPs by median, Fig. S3A), and ribosomes with several late E^OUT^ tRNA exit intermediates (combined 2.5% median, Fig. 2A, Fig. S3B, C).^61^ The peptidyl transfer-associated 70S A,P state was the most common ribosomal species in the cell (12.3% median, Fig. 2A), consistent with the abundance of late-decoding and early peptidyl transfer intermediates reported across species.^42–51^ E^OUT^ tRNAs exhibited anticodon loops shifted towards the head of the small subunit (Fig. S3D, E)^61^ and, similar to eukaryotic Z-tRNAs,^62^ occur prior to release of E-site tRNAs (Fig. 2A). The preceding E/E^OUT^ state is one where the E-site tRNA anticodon end is positioned between the E and E^OUT^ states (Fig. S3E). We also observed 70S complexes post-termination, in the canonical P/E pre-recycling state (Fig. S3F), and 50S complexes in a continuum of recycling states, as indicated by the presences of RRF and EF-G (Fig. 2A: magenta boxes, Fig. S3G-I).^2^

Recycling of the post-termination 70S occurs via the joint actions of RRF and EF-G, where GTP hydrolysis and the subsequent release of inorganic phosphate enable EF-G to break intersubunit bridges and split the (P/E-state) 70S into 50S and 30S.^63,64^ In the free 50S, these factors must then dissociate to allow for new translation. Still, little is known about the way this occurs. We resolved 50S complexes with both RRF and EF-G (6.3% median, Fig. S3G), 50S RRF complexes (8.7% median, Fig. S3H), and factorless 50S complexes (5.3% median, Fig. S3I) presumably competent for initiation, indicating that recycling factor dissociation occurs sequentially. However, >86% of 50S complexes (excluding putative biogenesis complexes described below) were observed with density corresponding to a tRNA in the E-site (Fig. S3G-I). Similar tRNAs were previously observed in vitro by time-resolved^64^ and ensemble^65,66^ single-particle cryo-EM analyses of *E. coli* ribosomes, but their physiological relevance has remained uncertain. 50S E-site tRNAs would be expected to negatively influence translation initiation by clashing with 30S-bound initiator tRNAs during subunit joining. Thus, E-site tRNAs likely dissociate during 70S formation, but the implications of this additional step in translation are not clear.

We additionally resolved putative biogenesis complexes (Fig. S3J-N, Table S1), which included a late biogenesis 30S (2.4% median, Fig. S3J) with weak or absent protein S3 density (Fig. S3K) and open head conformation (Fig. S3L), similar to previously-reported *E. coli* late biogenesis 30S structures.^67^ Likewise, we observed mid-biogenesis 50S-like complexes lacking substantial density in the peptidyl transferase center (PTC) (0.4% median, Fig. S3M), and late or inactivated^68^ 50S-like complexes with abnormal 23S rRNA helices 68 and 69 (2.3% median, Fig. S3N). Formation of the PTC and proper folding of these helices are implicated as important steps of convergence in large subunit biogenesis.^69^

Our analysis further showed that *M. pneumoniae* heavily depends on transcription-translation coupling during both translation initiation and elongation (Fig. 2A: dotted boxes, and detailed below). By median, 7.8% and 10.2% of all RNPs per cell were 30S subunits and 70S ribosomes coupled to RNAP, respectively. In *E. coli*, genes regulated by the alternative transcription-translation coupling factor RfaH are highly reduced in SD sequences,^70^ and it is proposed that the increased efficiency of transcription-coupled translation initiation is required for their effective translation.^71^ *M. pneumoniae*’s low occurrence of SD sequences^29^ may be compensated for in a similar fashion.

Finally, by analyzing the abundances of all classified RNPs as a function of cell size, we identified a different distribution of complexes in smaller cells (bottom 25% by volume) compared to medium-sized (middle 50%) or larger (top 25%) *M. pneumoniae* (Fig. 2B). We reasoned that smaller cells are those that have recently divided, in line with quantification of their total RNPs (Fig. 1G-J). Although a quiescent state cannot be fully excluded, we did not observe RNPs dimerized for hibernation, or with apparent hibernation factors,^72^ and the median volume of these cells (0.016 μm^3^) was substantially smaller than the measured median volume of stationary phase *M. pneumoniae* (0.075 μm^3^).^73^ Smaller cells, consistent with their reduced 70S/[50S+30S] ratios (Fig. 1I), had higher proportions of termination and recycling-related RNPs (Fig. 2B), including 70S complexes with P/E tRNAs (3.24-fold median), 30S complexes with IF3 (2.21-fold median), and 50S complexes with RRF and EF-G (1.43-fold median). Our results thus suggest that the transition between cell division and growth may involve change in the translation regulation program from termination and recycling to initiation and elongation.

### Translation initiation proceeds through a preferred order of factor association

To regulate translation initiation, IFs 1, 2, and 3 are thought to coordinate with the 30S, mRNA, and initiator tRNA for start codon recognition.^2,4^ The process is likened to a series of kinetic checkpoints where multiple independent steps gatekeep proper assembly, but it remains unclear whether initiation follows one or more pathways in cells.^3–5^ We therefore reconstructed an initiation pathway based on the assumption of sequential factor binding, and in line with the literature.^2–5,7,74^ Although our results are not dispositive of the concept that initiation factors generally bind without a strict assembly pathway,^75^ our analyses pointed to a preferred order in *M. pneumoniae* (Fig. 3A-G).

**Figure 3:**
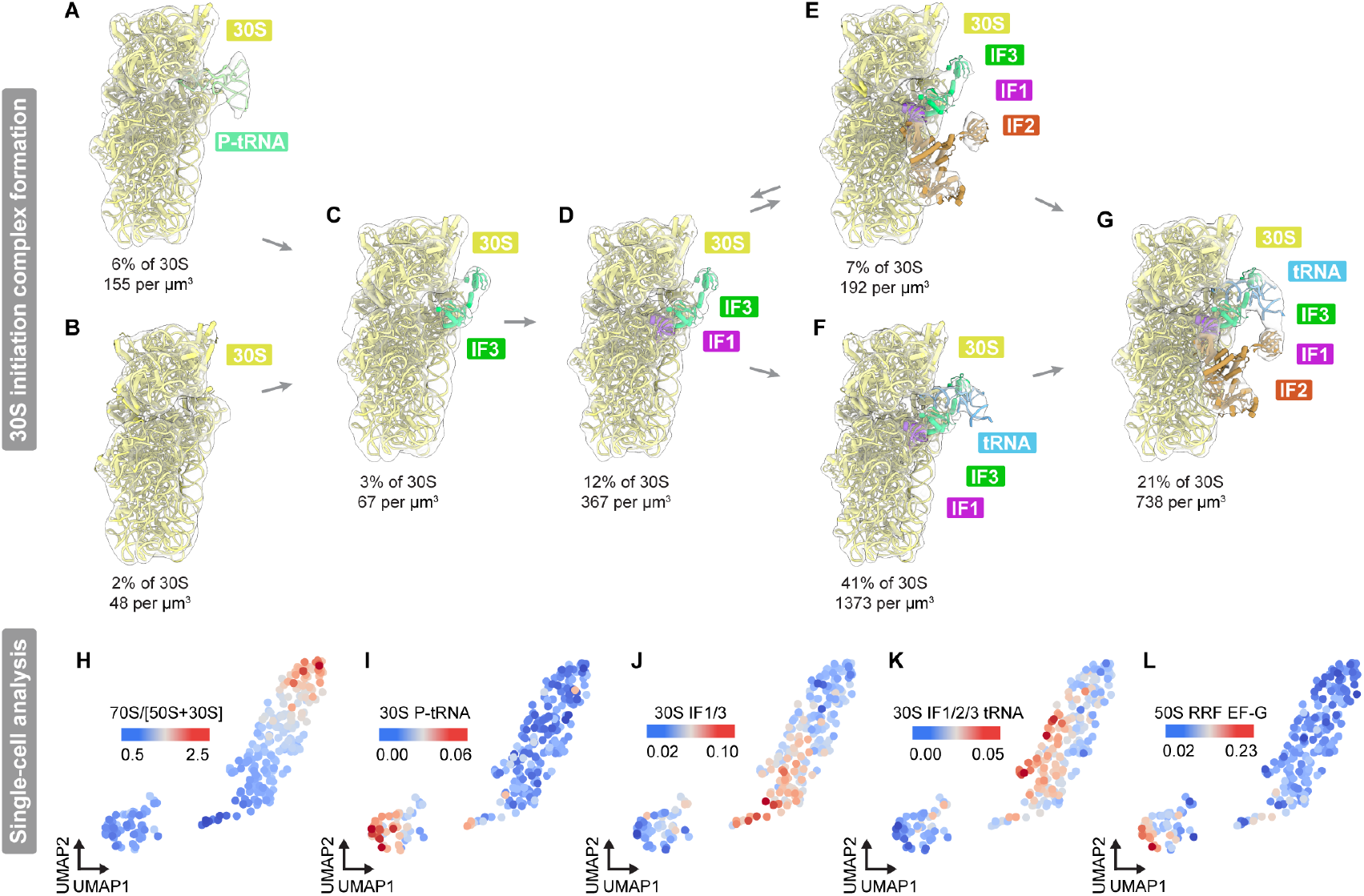
Translation initiation proceeds through a preferred order of factor association. **(A-G)** Reconstructions depict the buildup of factors during translation initiation. Maps (transparent) are fitted with molecular models (cartoon) derived from combinations of experimental (modified PDB 7OOC)^45^ and predicted models (Methods). **(A)** 30S with P-tRNA (9.4 Å). **(B)** Factorless 30S (18.3 Å). **(C)** IF3-bound 30S (11.2 Å). **(D)** IF1/3-bound 30S (7.5 Å). **(E)** IF1/2/3-bound 30S (9.2 Å). **(F)** IF1/3, initiator tRNA-bound 30S (5.6 Å). **(G)** IF1/2/3, initiator tRNA-bound 30S (6.6 Å). IF2 is represented in a stable form; the central GTPase domain was not resolved in the flexible form (Fig. S2C). **(H-L)** Single-cell analyses of translational complex abundance visualized by UMAP (Methods). Projections are coloured by the ratio of assembled 70S to the total of free 50S and 30S **(H)**, or by the per-cell RNP fractions of **(I)** 30S P-tRNA, **(J)** 30S IF1/3, **(K)** 30S IF1/2/3 tRNA, and **(L)** 50S RRF EF-G complexes. Dots represent individual cells (n=254). Related to Figures S2 and S3, Table S1.

It is noteworthy that 30S complexes with deacylated P-site tRNAs^73^ or with initiator tRNAs^74^ produce similar maps, where both tRNAs bind the 30S at the P site, making it difficult to differentiate them based on structure alone. Therefore, to place identified 30S P-tRNA complexes in the proposed initiation sequence, we clustered single cells using the uniform manifold approximation and projection (UMAP) algorithm (Methods),^76^ visualizing cells in a trajectory of translational activity via their 70S/[50S+30S] ratios (Fig. 3H). We found 30S P-tRNA complexes (Fig. 3I) to be enriched in different cell populations than 30S IF1/3 (Fig. 3J) and 30S IF1/2/3 tRNA complexes (Fig. 3K). Instead, cells high in 30S P-tRNA complexes clustered with cells enriched for recycling 50S RRF EF-G complexes (Fig. 3L), and both were coincident in cells with small volumes (Fig. S3O). These observations support an assignment of the 30S P-tRNA as a post-termination complex, and are consistent with IF3’s subsequent but early binding in initiation^3,77^ and long dwell time on the 30S.^77^

In our reconstruction of the initiation sequence, we therefore placed 30S P-tRNA complexes (Fig. 3A) alongside factorless 30S complexes (Fig. 3B). Both can then be bound by IF3 (Fig. 3C). In addition to its role in start codon recognition,^7^ IF3 is known to dissociate the deacylated P-tRNA^63,78,79^ and prevent improper reassociation of the 50S.^63,78^ Additional density in IF1’s position indicates its subsequent binding (Fig. 3D),^80^ followed by a possible split in the pathway. On one hand, we observed association of IF2, which is understood to help recruit the initiator tRNA and coordinate 50S binding,^81,82^ forming a 30S IF1/2/3 complex (Fig. 3E). On the other hand, we found the initiator tRNA substantially more often bound to the 30S without IF2 (Fig. 3F). By median, this 30S IF1/3/tRNA complex was the second most common RNP in the cell (10.8%, Fig. 2A). Finally, we resolved the 30S IF1/2/3/tRNA complex, which would be joined by the 50S once a start codon is recognized (Fig. 3G). All observed 30S structures in native cells, save biogenesis complexes (Fig. S3K), had open head conformations.^7^ In line with single-molecule experiments,^3,77^ our results support a model in which IF2 preferentially associates to the 30S after recruitment of the initiator tRNA. This interpretation is unexpected given IF2’s canonical role in initiator tRNA recruitment, but is consistent with the finding that IF2 binding does not appear to affect the positions of other IFs or initiator tRNA in *T. thermophilus* reconstitutions.^7^ Our data are not consistent with IF2 and the initiator tRNA binding simultaneously on the 30S.^74^ Indeed, kinetics experiments suggest the initiator tRNA can bind before IF2, depending on their relative concentrations and on the presence of other IFs,^74^ and a similar trajectory has been suggested based on analysis of ensemble and time-resolved cryo-EM studies.^4,7,79^ Furthermore, the high abundance of the 30S IF1/3/tRNA complex (41% of 30S) indicates that the transition to full 30S IF1/2/3/tRNA is an important checkpoint in the initiation pathway. Together, our results integrate single cell clustering and structural quantification to support a favoured pathway for translation initiation.

### Elongating transcription-translation complexes are coupled and collided in native cells

The lack of a nucleus in prokaryotic cells enables physical coupling of RNAP and the ribosome during gene expression^13–17,21–23,83^ and thus represents an additional level of cross-regulation. Structurally, this coupling is bridged by transcription factors, but the exact nature of the interaction varies across bacteria; *E. coli* uses NusG as the primary bridging factor,^16,17^ while *M. pneumoniae* uses NusA, leading to the emergence of different interfaces between RNAP and the ribosome.^20,83^ Functionally, the progress of an elongating ribosome along a nascent mRNA increases the processivity of RNAP (at the cost of fidelity), and reduces its rates of backtracking and premature termination.^13^ A trailing ribosome can also help to resolve RNAP stalls,^13,14,84^ and several structures of elongating transcriptiontranslation complexes (eT-TC) visualize the interaction between a stalled RNAP and colliding 70S.^15–17,21^ Yet, it remains unclear whether, or how often, this kind of collision occurs in cells under normal growth conditions.^14,17,18^

In line with our previous work,^20^ subclassification of 70S ribosomes revealed the presence of stably (Fig. 4A) or flexibly (Fig. 4B) associated eT-TCs in native cells. Unexpectedly, our expanded classification (Methods) now resolved collided eT-TCs (Fig. 4C, eT-TC comparison in Fig. 4D). We modeled the *M. pneumoniae* RNAP with AlphaFold3^85^ and rigid-body fitted the model into the maps (Fig. S4A-E, Methods).^86^

**Figure 4:**
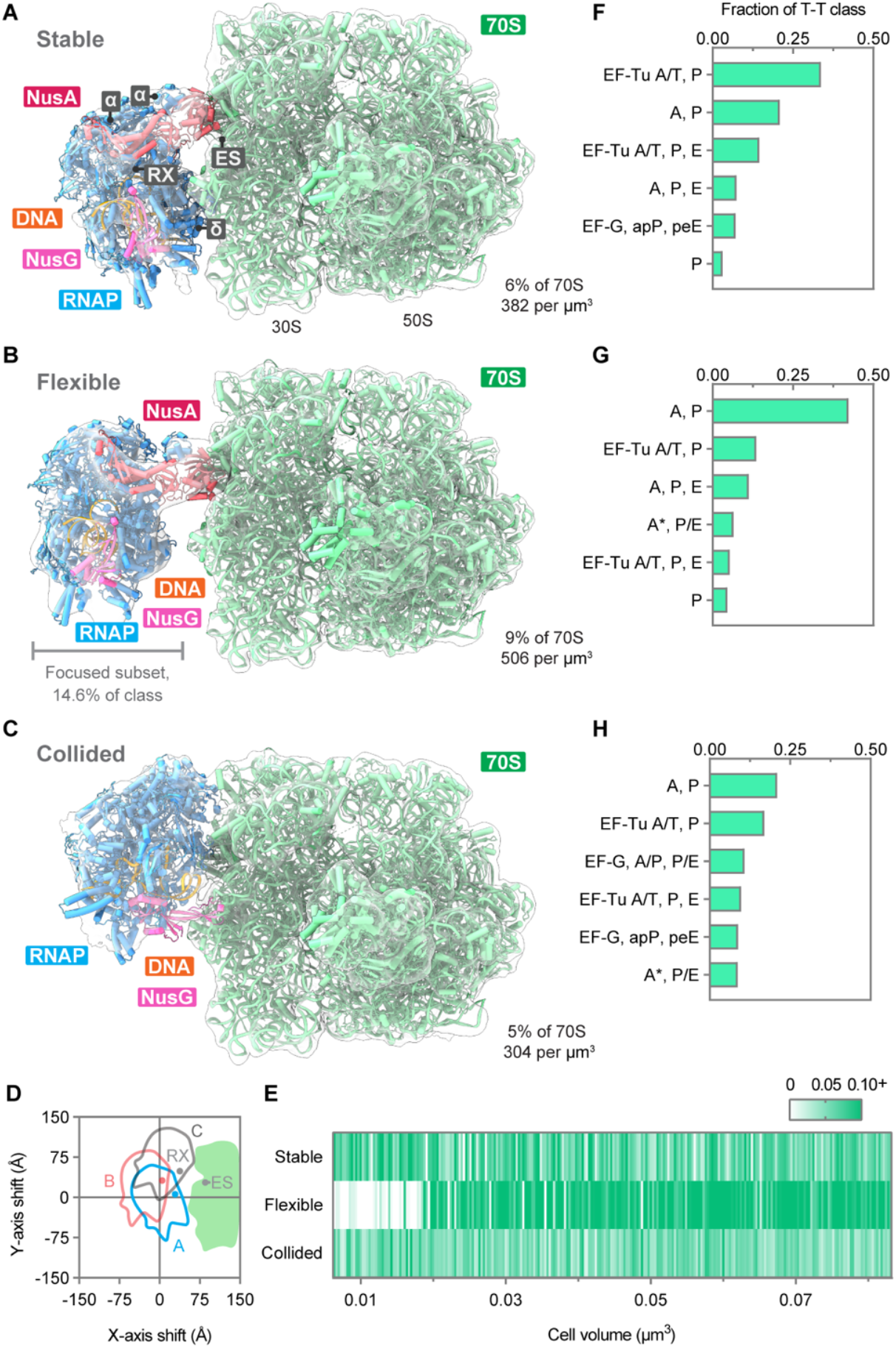
Elongating transcription-translation complexes are coupled and collided in native cells. **(A-C**) Maps of elongating transcription-translation complexes in native cells (transparent) fitted with molecular models (cartoon) derived from combinations of experimental (PDB 7PAI)^45^ and predicted models (Methods). **(A)** Stable eT-TC (6.5 Å). RX indicates the RNAP RNA exit channel, ES indicates the ribosomal mRNA entry site, *α* and *δ* indicate RNAP subunits. **(B)** Flexible eT-TC: composite map where RNAP was resolved with focused refinement of a subset of complexes and represents the highest likelihood conformation. The focused subset RNAP map (22.1 Å) is shown alongside the 70S consensus region (5.8 Å). **(C)** Collided eT-TC (7.2 Å). **(D)** Comparison of the relative RNAP positions in the (A) stable, (B) flexible, and (C) collided maps with respect to the 30S body. **(E)** Per-cell fraction of 70S complexes in each eT-TC conformation, ordered by cell volume (n=254 native cells). **(F-H)** Fractions of 70S with different tRNA and elongation factor states for **(F)** stable eT-TCs (n=3261 complexes), **(G)** flexible eT-TCs (n=4611), and **(H)** collided eT-TCs (n=2584). Related to Figures S2, S4.

The stable eT-TC (Fig. 4A) is bridged by NusA at the 30S mRNA entry site, as we previously reported,^20^ and the ordered N-terminus of RNAP’s *Firmicutes*-conserved additional δ subunit^87^ makes contact with the 30S body (Fig. S4F). Here, improved modeling of NusA identifies NusA’s interaction with RNAP to occur primarily through the two C-terminal domains of the RNAP α-dimer (Fig. 4A). Our analysis found that the stable eT-TC accounted for 6.3% of 70S complexes and 31% of all eT-TCs (Fig. S2A).

The flexibly associated form (Fig. 4B) was more common, and accounted for 9% of 70S complexes and 44% of all eT-TCs in native cells (Fig. S2A).

While our earlier estimates of this complex’s abundance were higher, we previously could not unambiguously resolve RNAP in this form.^20^ Indeed, single-particle cryo-EM studies of reconstituted T-TCs show similar flexibility and have required additional image processing to resolve a subset of RNAPs in any particular orientation.^13,16–18,21,22^ Here, we generated a composite map by recentering the flexible RNAP density and performing local classification, allowing us to visualize a subset of RNAPs in one high-occurrence conformation distal to the 70S (Fig. 4B, Methods). In contrast to the stable form, RNAP’s δ subunit does not contact the 30S and the interaction is bridged only by NusA and mRNA (the flexible transcript cannot itself be resolved). The loss of direct RNAP-30S interactions, and the architecture of this flexible complex, are analogous to reported structures of eT-TCs undergoing long-range coupling^88^ where the mRNA may loop out to substantial intervening lengths.^14^

The collided eT-TCs (Fig. 4C) accounted for 5% of 70S complexes and 25% of all eT-TCs in native cells (Fig. S2A). Similar to our previous collided structure^20^ in the presence of the RNAP-stalling antibiotic pseudouridimycin (PUM),^89^ NusA was excluded, and the body of the RNAP was rotated and shifted such that the RNA exit site was positioned at minimum distance on top of the 30S mRNA entry site (Fig. 4D). Furthermore, NusG (present in all eT-TCs forms) was tilted away from the DNA and appeared to contact the 30S body via its N-terminal domain. This complex is structurally consistent with collided *E. coli* eT-TCs (Fig. S4G), unlike the coupled eT-TCs forms (Fig. S4H).^15–17^ Finally, plotting the different eT-TC fractions across cells (Fig. 4E) showed that flexible eT-TCs were notably less common in smaller cells, while stable and collided eT-TCs were prevalent across all cell sizes. Experiments with *E. coli* reconstitutions suggest that flexibility in T-TCs enables mRNA accessibility via looping-out between RNAP and the ribosome, and thereby allows for regulation by other factors.^14,23,88^ Consistent with the changes to translation itself (Fig. 2B), regulation during T-T coupling may therefore occur differently in smaller compared to larger cells.

The ribosomes within stable eT-TCs were found primarily in the EF-Tu×A/T,P (33.7% of 70S) and A,P (20.9%) states (Fig. 4F),^45^ a departure from the total 70S population (EF-Tu×A/T,P: 17.0%, A,P: 35.0%). Flexible eT-TCs (Fig. 4H) were particularly enriched for A,P state ribosomes (42.0%). Ribosomes in native collided eT-TCs (Fig. 4H) had a broad distribution of EF and tRNA states, distinct from other eT-TCs, and in contrast to collided eT-TCs in PUM-treated cells which were primarily enriched for EF-G translocation intermediates.^20^ These data are consistent with the suggestion that during native collisions with RNAP translation elongation continues, but slows.^14^ As such, native collided eT-TCs are likely functional assemblies involved in resolving transiently stalled RNAPs.

While it has been suggested that collisions may be atypical or only occur when artificially induced,^17,18,14^ we found that collided eT-TCs were common in *M. pneumoniae* cells under normal growth conditions, in agreement with the essentially ubiquitous RNAP pause sites in prokaryotic DNA.^90–92^ Thus, our analyses resolve the question of whether collisions are relevant in the physiological context of a cell.

### Transcription and translation are physically linked during initiation

Interaction between RNAP and the ribosome can also occur prior to start codon recognition, during translation initiation, and recent studies have described the architectures of initiating transcription-translation complexes (iT-TCs) in different bacteria via in vitro reconstitutions.^19,21–23^ However, these studies highlight regulatory elements uncommon or absent in *M. pneumoniae*,^29,33^ suggest that RNAP can undergo extensive repositioning in the transition between coupled translation initiation and elongation, and do not address the prevalence of transcription-coupled translation initiation itself.

Our structural analysis revealed a substantial number of 30S subunits coupled to RNAP, in addition to the eT-TCs described above (Fig. 5A). We resolved iT-TCs in stable (Fig. 5B, 1.8% of 30S) and flexible forms (Fig. S4I, 28% of 30S), which both exhibited open 30S head conformations (Fig. S4J). Together, iT-TCs constituted 30% of 30S particles per cell (Fig. 5C). Similar in conformation to eT-TCs (Fig. S4K), iT-TCs included NusA and NusG and additionally translation initiation factors. We modeled IFs 1/3 and the initiator tRNA (Methods), although IF2 was also present in lower numbers in agreement with the proportions we quantified for the initiation sequence (Fig. 3A-G, Table S1). Unlike the *E. coli* or *T. thermophilus* reconstitutions,^19,21,22^ we did not resolve RNAP near the aSD. Instead, like eT-TCs, our iT-TC maps position RNAP closer to the 30S body and mRNA entry site (Fig. S4K, H). A similar state has been suggested as one of two potential pathways for RNA delivery in *E. coli*,^22^ although the RNAP-30S interface is distinct. Our data thus indicate that no major repositioning of RNAP is required in *M. pneumonaie* for the transition from coupled translation initiation to coupled translation elongation.

**Figure 5:**
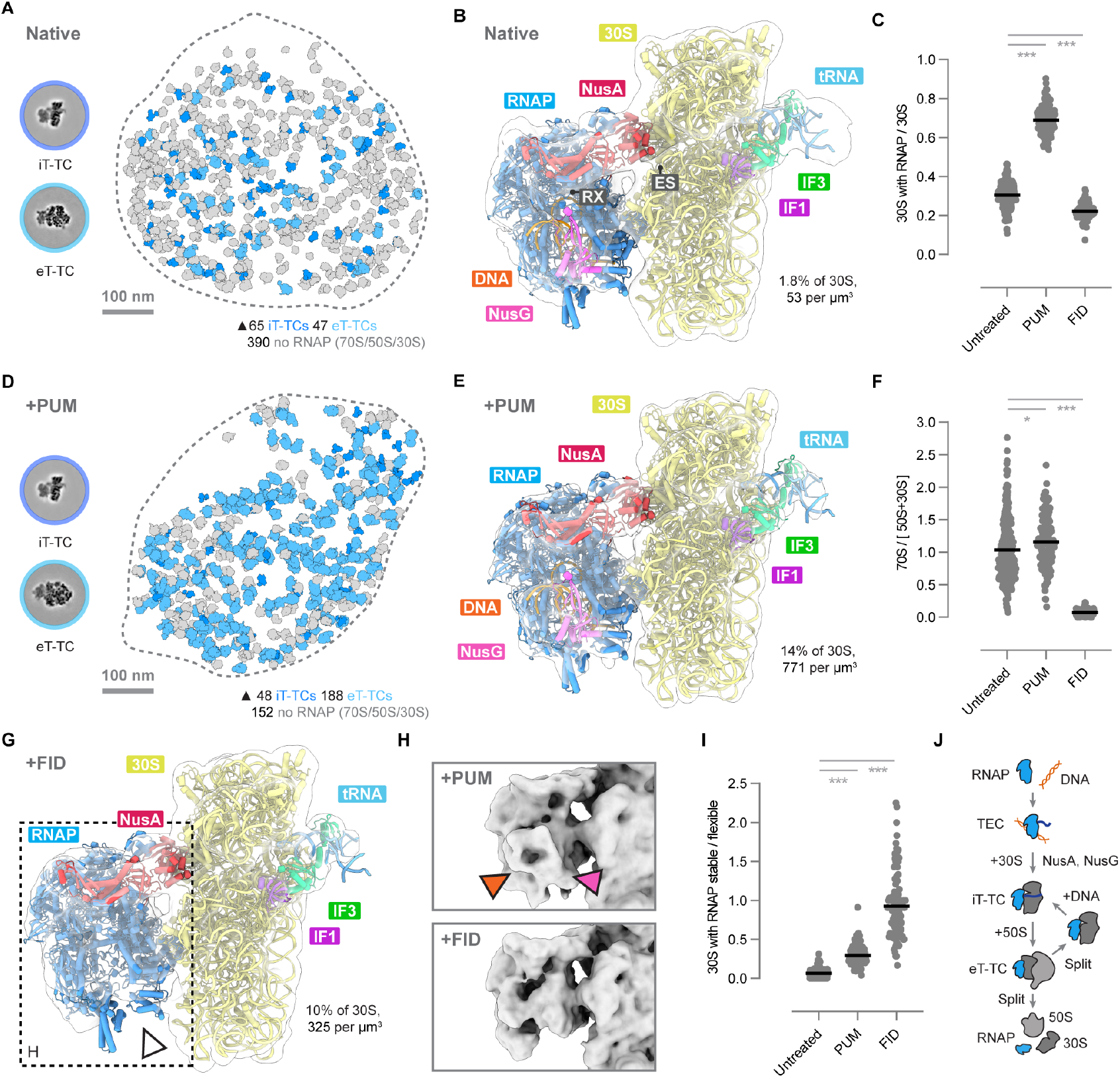
Transcription-translation linkage suggests a threading-based reinitiation mechanism. **(A)** Central slices through T-TCs reconstructions (left) corresponding to annotations of all T-TCs (blue) in a native cell (right; same cell as Fig. 1A). Slices were taken through stable form T-TCs for visualization. **(B)** Stable iT-TC in native cells (17.6 Å). RX indicates the RNAP RNA exit site, and ES indicates mRNA entry site. **(C)** Quantification of 30S complexes associated with RNAP in native untreated, PUM-treated, and FID-treated cells. **(D)** Central slices through T-TCs (left) corresponding to annotations of all T-TCs (blue) in a PUM-treated cell (right). Slices were taken through stable iT-TC and collided eT-TC for visualization. **(E)** Stable iT-TC in PUM-treated cells (10.2 Å). **(F)** The ratio of assembled 70S to free 50S + 30S subunits in native untreated, PUM-treated, and FID-treated cells. **(G)** Map of the stable iT-TC in FID-treated cells (9.5 Å). White arrowhead indicates the absence of density corresponding to the DNA and NusG regions of the map. **(H)** Inset view of RNAP in FID and PUM maps compared at the same density threshold. The orange arrowhead indicates the position of density corresponding to DNA, and the magenta arrowhead indicates the position of density corresponding to NusG. **(I)** The ratio of stable iT-TCs to flexible iT-TCs in native untreated, PUM-treated, and FID-treated cells. **(J)** A proposed model for translation reinitiation mediated by co-transcriptional threading of mRNA into the 30S. While 30S complexes are originally recruited to the transcription elongation complex (TEC), the machineries may remain associated post-termination to reinitiate coupled transcription and translation. Dots in plots represent individual cells and lines represent mean. Maps (transparent) fitted with molecular models (cartoon) were derived from rigid-body fitting of experimental (modified PDB 7OOC)^45^ and predicted models (Methods). Native (n=254), PUM-treated (n=124), FID-treated (n=100) cells. By Student’s T test: ns=not significant, *p < 0.05, **p < 0.01, ***p < 0.001. Related to Figures S2, S4, S5, S6, and Table S1.

To further investigate transcriptional coupling during translation initiation, we analyzed cells treated with PUM (Fig. 5C, D), introduced to inhibit transcription elongation.^89^ Following classification (Fig. S5), the resulting map was of sufficient resolution to model the architecture of the stable iT-TC (Fig. 5E). Consistent with our previous observation that PUM increases the abundance of eT-TCs,^20^ the treatment also substantially increased the total iT-TC fraction to 69% of 30S/cell (Fig. 5C). But unlike PUM eT-TCs, where the elongating 70S collides with the stalled RNAP (Fig. S4L), iT-TC maps in native and PUM-treated cells showed no apparent structural differences (Fig. S4M). We attribute this to the lack of force generated by the free 30S in contrast to elongating 70S complexes. The buildup of iT-TCs following PUM treatment (inhibiting transcription elongation) is consistent with single-molecule kinetics experiments suggesting that *E. coli* NusG assists the recruitment of 30S complexes to nascent mRNA,^93^ and sequencing experiments showing that *E. coli* NusA and NusG are recruited to RNAP only after transcription elongation begins.^94^

In summary, we find that physical transcription-translation coupling is an extensively used strategy for regulation during translation initiation in *M. pneumoniae*. Unlike the structures obtained from *E. coli* or *T. thermophilus* reconstitutions, the iT-TCs we resolved in *M. pneumoniae* do not rely on mRNA recruitment elements like the SD sequence or protein S1.^21,22^ Given *M. pneumoniae*’s low SD ORF fraction and lack of S1,^29,34^ noncanonical initiation is likely to be important in this bacterium and in other bacteria, or sequences, with similar characteristics.^24^

### Transcription-translation linkage suggests a threading-based reinitiation mechanism

The positioning of RNAP directly above the mRNA entry site in our iT-TC maps (Fig. 5B, E) suggested the unconventional possibility of direct mRNA threading into the 30S at the onset of translation initiation. To investigate this hypothesis, we treated cells with fidaxomicin (FID), which prevents transcription initiation by blocking new RNAP association with DNA,^95–97^ but allows ongoing transcription to continue and terminate. This resulted in a buildup of free ribosomal subunits and a decreased 70S/[50S+30S] ratio (Fig. 5F). Following classification (Fig. S6), we resolved both flexible and stable iT-TCs (Fig. 5G) accounting for 22% of 30S complexes, a decrease from 30% in native cells (Fig. 5C, Table 1). Stable iT-TCs in FID-treated cells (Fig. 5G) exhibited a similar conformation to untreated and PUM-treated stable iT-TCs but, crucially, FID iT-TCs lacked density in the RNAP DNA binding cleft and corresponding to NusG (Fig. 5H). These data indicate that ongoing transcription is not necessary to maintain the coupling of RNAP to 30S complexes, and that these assemblies may remain associated following transcription and translation termination. We also determined a substantial enrichment for stable complexes (Fig. 5I: untreated ratio 0.07, FID 0.92), suggesting a role for active transcription in the flexibility of iT-TCs.

Based on these results, we propose a model (Fig. 5G) in which the 30S and RNAP within eT-TCs may remain coupled following termination. Thus, upon reinitiation of transcription, RNAP may thread mRNA directly into the coupled 30S. Such a mechanism, rather than the assumed model of complete disassembly, could increase the 30S mRNA loading efficiency and may be important for translation of genes with atypical or noncanonical 5’ UTRs. Yet, this raises the question of how exactly RNAP can dissociate from the ribosome. Appealingly, following ribosome assembly and eT-TC formation, we observed that collided complexes have lost the NusA bridge (Fig. 4C), suggesting that disassembly of T-TCs may be controlled by collision events, and consistent with recent single-molecule experiments^14^ demonstrating that transcription-translation coupling becomes short-lived in response to collisions.

### Translation regulation at the cell membrane

Ribosomes involved in co-translational translocation are subject to additional regulation by recruitment to the cell membrane, and through interactions with the translocation machinery. In addition to the cytosolic complexes discussed above, we found *M. pneumoniae* cells to contain a variety of membrane-associated 70S complexes (Fig. 6A-C) and a surprisingly large fraction of previously undescribed membrane-associated 50S subunits (Fig. 6D). Membrane-bound 70S ribosomes had elongation factor and tRNA occupancies (Fig. 6E) consistent with the overall translating population (Fig. 2A), indicating ongoing translation.

**Figure 6:**
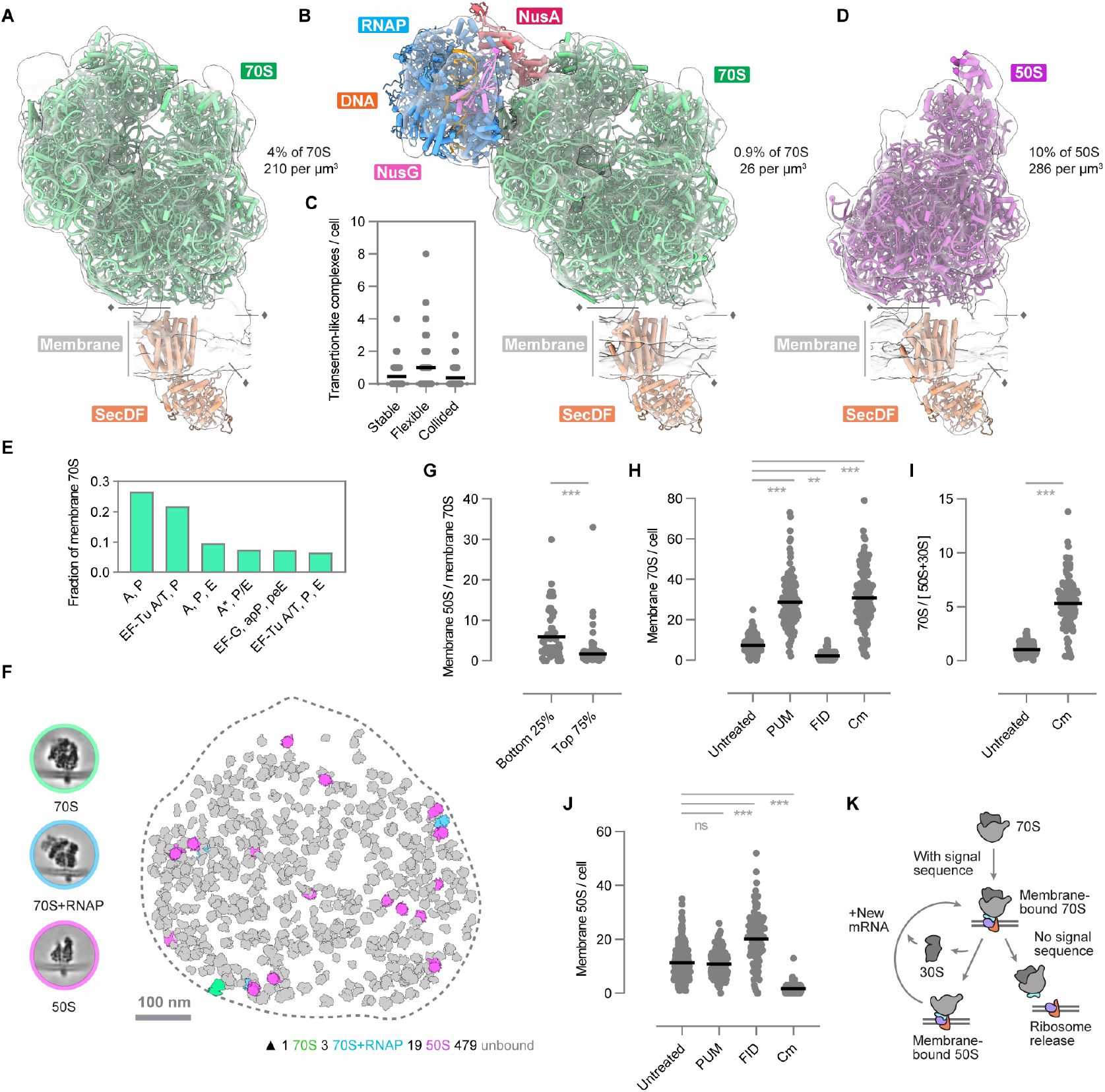
Translation complexes at the cell membrane crosstalk with the transcription and membrane-attachment machineries. **(A-D)** Diverse RNPs associated with the cell membrane. **(A)** Map of a membrane-bound 70S complex (transparent, 8.6 Å) fitted with a molecular model including SecDF derived from rigid-body fitting of experimental (PDB 7PAI)^45^ and predicted components (Methods). **(B)** Map of a transertion-like supercomplex (transparent, 26.7 Å) fitted with a molecular model including 70S (PDB 7PAI)^45^, SecDF, and RNAP in stable conformation (Methods). Complexes in (B) are a subset of (A). **(C)** Quantification of transertion-like supercomplexes per native cell classified by RNAP conformation. **(D)** Map of a membrane-bound 50S complex (transparent, 7.7 Å) fitted with a model of 50S (PDB 7PAT)^45^ and SecDF (Methods). Diamond motifs indicate unassigned density (detailed in Fig. S7). **(E)** Fractions of tRNA and EF states in membrane-associated 70S complexes. **(F)** Central slices through reconstructions (left) and corresponding annotations of all membrane-bound RNPs in a native cell (right; same cell as Fig. 1A). **(G)** The ratio of membrane-bound 50S complexes to membrane-bound 70S complexes in the smallest 25% (n=64) and largest 75% (n=190) of cells by volume. **(H)** Quantification of membrane-bound 70S complexes per cell in native, PUM-treated, FID-treated, and Cm-treated cells. **(I)** The ratio of assembled 70S complexes to free 50S + 30S subunits in native and Cm-treated cells. **(J)** Quantification of membrane-bound 50S complexes per cell in native, PUM-treated, FID-treated, and Cm-treated cells. **(K)** A proposed model for regulation of ribosome localization at the membrane. Dots in plots represent individual cells and lines represent mean. Native (n=254), PUM-treated (n=124), FID-treated (n=100), Cm-treated (n=121) cells. By Student’s T test: ns=not significant, *p < 0.05, **p < 0.01, ***p < 0.001. Related to Figure S5, S6, S7, S8, and Table S1.

Alongside these membrane-associated 70S ribosomes (Fig. 6A, 7.5/cell), we resolved transertion-like supercomplexes (Fig. 6B, 2/cell). The long-hypothesized structures^98–102^ are composed of transcribing RNAPs coupled to translating membrane-attached ribosomes. RNAPs in these complexes were associated in stable (0.5/cell), flexible (1/cell), and collided (0.5/cell) forms (Fig. 6C), similar to the relative distribution of eT-TC conformations in the cytosol. Bacteria may use such supercomplexes to expand the accessibility of the chromosome by physically tethering DNA to the cell membrane and organizing the structure of the nucleoid (if present) for localized protein synthesis.^103–106^ The presence of these supercomplexes has been suggested by indirect data over many years,^98–101,103–107^ and our maps provide the first structural evidence of their existence.

All membrane-associated translation complexes exhibited similar membrane-proximal densities (Fig. S7A, B). The shape of a transmembrane density protruding to the extracellular side suggested the presence of SecDF (108 kDa), a proton-conducing membrane chaperone associated with ATP-independent protein translocation.^108^ Indeed, fitting of a SecDF structure prediction explained the majority of extracellular and transmembrane density in the map (Fig. S7C, D).^85,109^ We speculated that the remaining transmembrane density may accommodate the translocon channel protein SecY (52 kDa), likely in complex with SecE (14 kDa) and SecG (8 kDa).^36,40,110,111^ However, the size of the density was insufficient to fit a predicted model of SecYEG (Fig. S7E, F). Furthermore, despite the limited local map resolution (due to severe preferential orientation, Fig. S7H), the shape and position of the density were inconsistent with an experimental model of SecYEG in complex with the *E. coli* ribosome (Fig. S7G).^39^ To explore which proteins may explain the remaining density in the maps, we predicted structures of *M. pneumoniae* proteins known to be associated with membrane translocation with AlphaFold3,^85^ and evaluated their fits (Fig. S7I-P, Methods). The intracellular density was too small to fit the ATPase SecA (92 kDa), but matched previously reported positions of processing enzymes peptide deformylase (PDF, 25 kDa, Fig. S7I, J) and methionine aminopeptidase (MetAP, 28 kDa, Fig. S7K,L).^112–114^ Rigid-body fitting showed that each protein could be accommodated, but the limited local map resolution precluded unambiguous assignment. We note that *M. pneumoniae* PDF appears to lack the conserved helix by which *E. coli* PDF binds the ribosome.^113^ The intracellular density could similarly accommodate a model of the signal recognition particle receptor FtsY (39 kDa), or SRP itself (50 kDa, Fig. S7M,N). However, the fits remained ambiguous, and these proteins are known to associate with the ribosome in many positions and orientations.^40^ Our final candidate was YidC (44 KDa), an integral membrane protein capable of either SecYEG-dependent^115^ or independent membrane protein insertion, and known to form a complex with SecYEG, SecDF and YajC.^116,117^ *M. pneumoniae* YidC (Fig. S7O) lacks the extracellular domain present in *E. coli*,^115^ but fit with the highest cross-correlation values of all candidates (Fig. S7O,P). Given the uncertainty in assignment, we only included SecDF in our molecular models for the transmembrane region and could not account for functional translocation activity despite the apparent active translation (Fig. 6E). The lack of density consistent with SecYEG suggests *M. pneumoniae* may have developed noncanonical translocation strategies during its reductive evolution, potentially making use of SecDF itself for SecYEG-independent translocation. To support a departure from the canonical mechanism, we obtained an in-cell map of the *E. coli* 70S anchored to the membrane (Fig. S7Q-T) where, while preferential orientation posed a similar issue to that in our *M. pneumoniae* data, transmembrane density consistent with SecYEG was evident (Fig S7R, S). Furthermore, the angle at which the *E. coli* ribosome was attached to the membrane was substantially different to the *M. pneumoniae* ribosome (8° rotation, Fig. S7U), consistent with the hypothesis that co-translational translocation occurs differently in these bacteria. Our recent work detailing a *Mycoplasma*-specific translocation-folding complex,^41^ which includes both SecDF and SecYEG, further supports the possibility that these minimal bacteria have evolved unusual translocation strategies.

### Large subunits remain membrane-anchored following translation termination

Our analyses identified a substantially larger number of membrane-bound 50S subunits (Fig. 6D, 11/cell) in comparison to membrane-bound 70S ribosomes in native cells (Fig. 6A, 7.5/cell). Visualization of all membrane-associated RNPs underscores this disparity in abundance (Fig. 6F): we found that the ratio between membrane-bound 50S and 70S complexes is significantly higher in smaller cells than in larger ones (Fig. 6G, means: 5.9x vs. 1.7x), consistent with the lower 70S/[50S+30S] ratio in smaller cells (Fig. 2B). These observations led us to hypothesize that the large subunit could be stably bound to the membrane, potentially only dissociating during translation of a cytosolic protein, as suggested to occur at the endoplasmic reticulum (ER) in mammalian cells.^118,119^

To investigate this possibility, we analyzed the abundances of membrane-associated complexes following targeted antibiotic perturbations. We first perturbed protein synthesis. Treatment with PUM to stall transcription elongation, or with the translation elongation inhibitor chloramphenicol (Cm, Fig. S8),^120^ both resulted in significantly increased counts of membrane-bound 70S complexes (Fig. 6H, PUM: 29/cell, Cm: 31/cell). Both Cm and PUM increased the 70S/[50S+30S] ratio (Fig. 6I, Fig. 5F), and Cm also reduced the number of membrane-bound 50S complexes (Fig. 6J, Cm: 2/cell). These perturbations resulted in stalled translation elongation, preventing recycling of 70S ribosomes and translation reinitiation, thus accumulating 70S complexes at the membrane. We next blocked the initiation of gene expression. Treatment with FID substantially decreased the 70S/[50S+30S] ratio (Fig. 5F) and led to a reduction in membrane-bound 70S complexes as gene expression completed (Fig. 6H, 2/cell). Due to post-termination 70S recycling, this resulted in a significant accumulation of membrane-bound 50S complexes (Fig. 6J, 20/cell). Consistent with our hypothesis, these data show that when reinitiation is inhibited, recycled large subunits remain membrane-bound. Our results therefore support a model where after initial ribosome recruitment to the membrane, elongation, and termination, the large subunit’s membrane attachment is regulated by the reinitiation of translation (Fig. 6K).

Since a similar mechanism was originally suggested in a mammalian system,^118,119^ we reanalyzed the EMPIAR-11751 cryo-ET dataset^50^ and localized rare large subunits present in the data to derive a map of the human 60S large subunit bound to ER-derived membranes via the SEC61-TRAP-OSTA translocon (Fig S7V, W). Thus, our results suggest that reinitiation-based control of membrane attachment, originally proposed to occur in mammalian cells but not resolved structurally before this work, is likely to be generally conserved.^118,119^

## Conclusions

Using in-cell cryo-ET to resolve complexes associated with the translation initiation, elongation, termination, and recycling phases, and their structural coordination with parallel processes in *M. pneumoniae*, our analysis paves the way for system-wide investigations of the molecular landscape of gene expression within cells under normal growth conditions and in those subjected to perturbations. We combined characterization of cellular properties with structural analyses to uncover and quantify regulatory interactions that modulate the translation cycle. Our integration of single-cell analysis and computational classifications provides the first in-cell structural quantification of translation initiation dynamics, and our investigation of transcription-translation coupling resolves the question of whether collisions are physiologically relevant under normal conditions.^14,17,18^ Our data quantify the extent of transcription-translation coupling during initiation, where the in-cell *M. pneumoniae* RNAP-30S interface appears distinct from structures derived from *E. coli* and *T. thermophilus* reconstitutions.^13,16,17,21–23^ Furthermore, we suggest an unexpected role for transcription-translation coupling in reinitiation. Our analysis of membrane-associated ribosomes visualizes a long-hypothesized gene expression supercomplex that coordinates transcription, translation and membrane attachment,^98–100^ and by localizing membrane-bound large ribosomal subunits,^118,119^ suggests that the principle underlying control of ribosomal membrane attachment is conserved from minimal bacteria to human cells.

## Acknowledgements

We thank Mahamid group members for useful discussions, and especially L. Xue for generating data included in this work and preliminary analysis. For technical support we are grateful to E. Zagoriy, J. Bartho and S. Unger from the EMBL cryo-EM platform, and EMBL IT especially T. Hoffmann. For critical feedback on the manuscript we are grateful to O. Duss, N. Qureshi and L. Xue. R.K.J. acknowledges funding from the Independent Research Fund Denmark (0106-00010A). J.M. acknowledges funding from the EMBL and a Chan Zuckerberg Initiative grant for visual proteomics (2021-234620). This paper was typeset with the bioRxiv word template by @Chrelli: www.github.com/chrelli/bioRxiv-word-template.

## Author contributions

J.M.D. and J.M. conceptualized the study. J.M.D. and R.K.J collected data. J.M.D. performed analysis. J.M.D. and J.M. wrote the manuscript with input from all authors.

## Data, tool, and code availability

- Public repository data from Xue et al.,^45^ Fromm et al.,^121^ Zhang et al.,^61^ Jomaa et al.,^39^ Gemmer et al.,^50^ Natchiar et al.,^122^ and Xue et al.,^123^ were used.
- Softwares including SerialEM,^122^ SPACEtomo,^123^ Warp,^124^ AreTomo2,^125^ IMOD,^126^ PyTom,^127^ DeePiCt,^128^ EMAN2,^129^ RELION4,^130^ ChimeraX,^86^ ArtiaX,^131^ M,^132^ AlphaFold3,^85^ WarpTools,^133^ starparser,^134^ tomoDRGN,^109^ and GraphPad Prism were used.
- 140 cryo-ET maps have been deposited in the Electron Microscopy Data Bank (EMDB)^135^ and 17 models have been deposited in the Protein Data Bank (PDB).^136^ Accession numbers will be provided with publication.
- Structures used to create models, or for comparison, were obtained from the PDB or were generated with AlphaFold3.^84^
- Python scripts for polysome analysis, volume calculations, and UMAP plots are available at https://github.com/jmdobbs/cell_analysis_scripts

## Materials and Methods

### Experimental models

*Mycoplasma pneumoniae* M129-B7 cells were kindly provided by Prof. Jörg Stülke, and *Escherichia coli* BW25113 cells were kindly provided by Dr. Athanasios Typas.

### *M. pneumoniae* culture, vitrification, and sample preparation

Samples were prepared for cryo-ET experiments as previously described.^20^ Briefly, wild-type *M. pneumoniae* strain M129-B7 were maintained at 37° C in modified Hayflick medium containing 18.4 g/L Difco PPLO (Becton Dickinson), 100mM HEPES pH 7.4 (Biomol), 0.002% phenol red (Reagecon), plus 20% (v/v) Gibco horse serum (Thermo Fischer Scientific), 1% (w/w) glucose (Merck Millipore), and 1000 U/ml penicillin G (Sigma-Aldrich) added after autoclave cooling.^138^ Quantifoil R2/1 200 mesh gold grids (Quantifoil Micro Tools) were either plasma cleaned for 1 minute with a 90:10 argon:oxygen mixture (Fischione M1070), or glow discharged for 45 seconds (PELCO easiGlow), then UV irradiated for 30 minutes in a sterile laminar hood prior to seeding in 35 millimeter culture dishes. Vitrification was performed 16-20 hours after seeding to ensure cells were in growth phase. Vitrification was conducted with a manual plunger (manufactured by the Max Planck Institute of Biochemistry) or a Leica GP1 plunger (Leica Microsystems, 37° C, 40% humidity). Grids with adherent *M. pneumoniae* cells were briefly washed with phosphate buffered saline (PBS) containing 10 nm protein A-conjugated gold beads (Aurion), blotted from the back for 0.5-2 seconds, and plunged into a liquid ethane/propane mixture. For antibiotic treatments, compounds were added to the culture medium at 37° C at the following concentrations and for the specified amount of time before vitrification: PUM: cells treated at a concentration of 0.4 mg/ml for 15 minutes. FID: cells treated at a concentration of 0.4 mg/ml for 30 minutes. A 60-minute FID treatment was also performed, but single cells largely appeared lysed or had dissociated from the grid. Cm: cells treated at a concentration of 0.2 mg/ml for 15 minutes.

### Tilt series acquisition

Tilt series were collected on a Titan Krios G3i (Thermo Fischer Scientific) transmission electron microscope operated at 300 keV equipped with post-column energy filter (Gatan) and K2 or K3 direct electron detectors (Gatan) using SerialEM software.^124^ Tomograms were collected from -60° to +60° at 3° increment using the dose-symmetric tilt scheme^140^ and the following parameters: total fluence ∼120 e^-^/Å^2^, pixel size 1.71 Å (K2) and 1.631 Å (K3) or 1.329 Å (K3, Cm data only), defocus range -2 to -5 µm, nominal magnification 81000x and 53000x or 64000x, respectively. 70/254 untreated tomograms were acquired using a Volta phase plate^141^ in addition to the above parameters. Prioritizing completeness of annotation over maximum data size, analysis was performed on 254 tomograms of untreated cells, 124 tomograms of cells treated with PUM, 100 tomograms of cells treated with FID, and 121 tomograms of cells treated with Cm. Data collection parameters are provided in Supplementary Table 1.

### Tilt series processing, particle picking, and subtomogram analysis

Tilt series were motion- and CTF-corrected in Warp v1.09^126^ or a pre-release version of WarpTools,^135^ aligned with AreTomo2^127^ or IMOD Etomo,^128^ and reimported into Warp (or WarpTools) for tomogram reconstruction. Tomograms were reconstructed at 10 Å per pixel (Å/px) for visualization and particle picking.

Initially, 70S, 50S, and 30S references for template matching were generated from the data using manual picking in EMAN2,^131^ followed by classification and averaging of approximately 600 subtomograms per target in RELION v4.0.1 in single-particle mode.^132^ Template matching in PyTom v1.1^129^ was performed using these references low-pass filtered to 20 Å/px, followed by RELION classification and averaging.^132^ DeePiCt^130^ models were trained on these validated template matching picks and used to predict additional putative complexes. EMAN2 was used to manually validate the picks, exclude complexes outside of the target cell (if there was more than one cell in the tomogram), and manually pick additional potential complexes of interest (Fig. S1A). This exhaustive process was necessary to obtain essentially complete picking of all putative translation complexes.

To obtain consensus maps for the 70S, 50S, and 30S complexes, subtomograms were reconstructed at 5 Å/px in Warp at all putative positions and initially classified in RELION with 30S, 50S and 70S references. Classified particles of each consensus type were refined (Fig. S1A), and duplicate or overlapping particles were removed with a combination of starparser^136^ and RELION functions (overlap criterion: <100 Å). Local angular refinement was subsequently performed in RELION, and the resulting averages were used as references for M-correction of deformations and shifts in tilt series.^134,135^ Subtomograms at the original coordinates of all potential complexes were then reconstructed again, classified, and refined. M-correction was then performed in multi-species mode. This process was repeated 3 times in total, after which there was no apparent improvement in the number and completeness of classified complexes (Fig. S1A). M-correction was performed separately for each dataset due to different camera geometry or additional use of the Volta phase plate (which requires correction for phase shift). The datasets were then merged (Fig. S1A), combined averages were subject to local angular refinement in RELION, imported into M and subjected to pose refinement, exported as image stacks for tomoDRGN^109^ cleaning of persistent junk particles, again re-imported into M, and subjected to final M pose refinement with unbinned pixels (K2: 1.71 Å/px, K3: 1.631 Å/px). This process resulted in a 3.8 Å global resolution average of the 70S ribosome, a 4.2 Å average of the 50S large subunit, and a 4.8 Å average of the 30S small subunit (Fig. S1A, E).

Extensive classification then was performed at 5 Å/px in RELION using subtomograms from each of the three consensus reconstructions, as detailed in Figure S2. As in our previous work,^45^ in order to reduce variance, classification was performed in triplicate, the middle or most consistent number of particles was selected, and output classes were subjected to additional classification following a round of local refinement (with a spherical mask) as quality control for misclassification. Classification here was performed without alignments, restricting the resolution of the RELION expectation step to a maximum of 20 Å, and (generally) with spherical or ellipsoid masks around regions of interest (Fig. S2A). Multiple trials were performed to select optimal masking parameters and mask positioning. For flexible T-TCs, particles were recentered on RNAP and classification was performed with 7.5° local alignments and a spherical mask. The classification procedure described here enabled us to recover multiple additional classes not observed in the previous *M. pneumoniae* studies.^20,45^ In order to maintain consistency between datasets and accurately classify the effects of perturbations, the same procedure, including the same masks and references, was used for all classifications across treatments.

Once classified, local angular refinement was performed in RELION and complexes were re-imported into M for 3 rounds of final pose refinement at 2 Å/px, where final resolutions and Fourier Shell Correlation (FSC) curves were calculated for all identified complexes (reported in Fig. S2). For some rare low-resolution maps (*e*.*g*. recentered flexible T-TCs), complexes were re-imported into M at 5 Å/px. Mapping of complexes into tomograms was visualized with the ChimeraX plugin ArtiaX.^133^ The same procedures were performed for antibiotic-treated data originally collected at 1.631 Å/px (PUM and FID data, Fig. S5 and S6), or 1.329 Å/px (Cm data, Fig. S8).

For the human membrane-bound 60S, EMPIAR-11751 (which contains 893 tomograms of ER-derived vesicles from HEK293F cells)^50^ was reanalyzed using essentially the same procedure as described above. Models from PDBs 6QZP and 8B6L were rigid-body fitted into the map (Fig. S7V).^50,122^

### *E. coli* cryo-ET sample preparation, acquisition, and analysis

To obtain a map of the membrane-bound *E. coli* 70S, 296 tilt series were collected on 21 cryo-focused ion beam milled lamellae of native *E. coli* BW25113 cells. In brief, 1 milliliter of cell suspension was grown at 37° C until OD 0.6 in lysogeny broth (LB) medium with 1mM CaCl_2_ and 1 mM MgCl_2_, concentrated by centrifugation at 5000g for 1 minute, and resuspended in 16 µl medium (to OD ∼36). 4 µl suspension was applied to plasma cleaned (Fischione M170) Quantifoil R1/4 carbon-coated copper grids (Quantifoil Micro Tools), and samples were vitrified with a GP1 plunge freezing device (Leica Microsystems, 37° C, 40% humidity). Lamellae were milled to approximately 160 nm thickness at 12° milling angle using an Aquilos microscope (Thermo Fisher Scientific) with autoTEM software. These were prepared with gallium ions in a stepwise manner and currents gradually reducing from 1 nA (for rough milling) to 50 pA (for final polishing). Platinum was deposited before (sputtered and organometallic platinum) and after (sputtered only) milling.

Tilt series were collected on a Titan Krios G4 equipped with a SelectrisX energy filter and a Falcon4i camera (Thermo Fisher Scientific) using SerialEM^124^ and SPACEtomo^125^ softwares. Tomograms were collected from - 66° to +48° at 3° increment and -12° pretilt using the dose-symmetric tilt scheme^140^ and the following parameters: total fluence ∼120 e^-^/Å^2^, pixel size 1.93 Å, defocus range -2.5 to -5 µm, nominal magnification 64000x. Data were processed and analyzed using essentially the same strategy as described above, and models from PDB 8B0X^121^ and PDB 5GAE^39^ were rigid-body fitted into the map (Fig. S7R,S). Cell membranes (Fig. S7Q) were segmented with membrain-seg.^142^

### Molecular modeling

Consensus ribosomal structures were modeled with PDB 7PAI (70S), PDB 7PAT (50S) or 7PAU (50S with RRF), and PDB 7OOC (30S) from our previous work.^45^ For the 30S, the head region was separated from the elongation-stage model and rigid-body fitted into the (initiation-stage) untreated consensus 30S map with ChimeraX^86^ to better model the open initiation head conformation.

For initiation and recycling complexes, protein sequences were obtained from Uniprot^143^ and the above experimentally-derived models were combined via rigid-body fitting with AlphaFold3^85^ predictions of *M. pneumoniae* IF1 (Uniprot Q50298), IF2 (Uniprot P75590), IF3 (Uniprot P78024), EF-G (Uniprot P75544), or L10 and the L7/L12 stalk (Uniprot P75240 and P75239). tRNAs (E-tRNA on the 50S, P-tRNA on the 30S, initiator tRNA on the 30S) were modelled with a P-tRNA from PDB 7PAL.^45^

For ribosome-RNAP complexes, *M. pneumoniae* RNAP as a whole (α, α, β, β’, δ: Uniprot IDs Q50295, Q50295, P78013, P75271, P75090) was predicted with a 38 base-pair sequence of DNA adapted from one used in structural studies of the *E. coli* RNAP.^144^ The RNAP model (Fig. S4A, B) was predicted to be of good quality (Fig. S4B, C), especially in the RNAP core. Separate predictions of the RNAP core with NusG (Fig. S4D, Uniprot P75049) and NusA (Fig. S4E, Uniprot P75591) positioned the factors in orientations that matched the map density, and that partially matched our previous integrated model (Fig. S4F).^20^ The RNAP core of this previous model also closely matched the RNAP core predicted here. A combined model was constructed by merging the RNAP core, DNA, and transcription factor predictions in ChimeraX, and rigid-body fitting of the model into the stable eT-TC map was performed. Slight manual repositioning of the NusA N-terminal and KH domains was also performed with ChimeraX to better fit the maps. AlphaFold3 did not initially predict the interaction of the second RNAPα C-terminal domain with NusA’s KH domain, as it did for the first RNAPα C-terminal domain. Therefore, these elements were separately predicted and recombined into the full model via rigid-body fitting to the map. The flexible C-termini of the RNAPδ subunit and NusG were deleted from the model as they could not be resolved in the map. For the flexible eT-TC and iT-TC, RNAP-NusA-NusG was rigid-body fitted into recentered subset RNAP maps.

For the membrane-bound ribosome complexes, consensus 70S (PDB 7PAI), 70S with stable RNAP (described above), and 50S (PDB 7PAT) models were rigid-body fitted into maps. AlphaFold3 predictions of the translocation-associated components including *M. pneumoniae* SecDF (Uniprot P75387), SecY (Uniprot P75112) + SecE (Uniprot P75048) + SecG (Uniprot Q9EXD0), YidC (Uniprot Q59548), FtsY (Uniprot P75362), SRP (Uniprot P75054), PDF (Uniprot P75527), and MetAP (Uniprot Q11132) were obtained, and these were rigid-body fitted into maps (Fig. S7A-N). Other assessed predictions included most transmembrane-annotated *M. pneumoniae* proteins in Uniprot over approximately 25 kDa. Save for SecDF, local resolution was largely insufficient to unambiguously place or identify transmembrane components of the maps (Fig. S7E).

### Analysis of cellular properties

Cell volumes, polysome counts, and UMAPs were computed using custom python scripts. Briefly, cell volumes were computed by determining the convex hulls around coordinates of all putative complexes in manually annotated cells, and converting the resulting volumes from pixels to cubic micrometers. Polysome counts were determined by spatial distance analysis of 70S ribosome mRNA exit sites to mRNA entry sites for the nearest neighbor 70S ribosome: a histogram of distances was computed (Fig. S1E) and a conservative 7 nm cutoff, similar to our previous work,^45^ was selected to describe ribosomes in polysome chains. UMAP projections were computed using parameters including cell volumes, counts of 70S, 50S, and 30S complexes per-cell, and fractions of total RNPs for each identified complex.

### Quantification and statistical analysis

Plots were produced in GraphPad Prism where statistical analysis was also performed. In all cases, unpaired Student’s T tests assuming non-equal standard deviations were used to compute the statistical significance of differences. R^2^ was used to describe goodness of fit for linear regressions and RMSE was used to describe goodness of fit for nonlinear regressions. Detailed information about quantification and statistical testing is included in the relevant figure legends.

**Figure S1:**
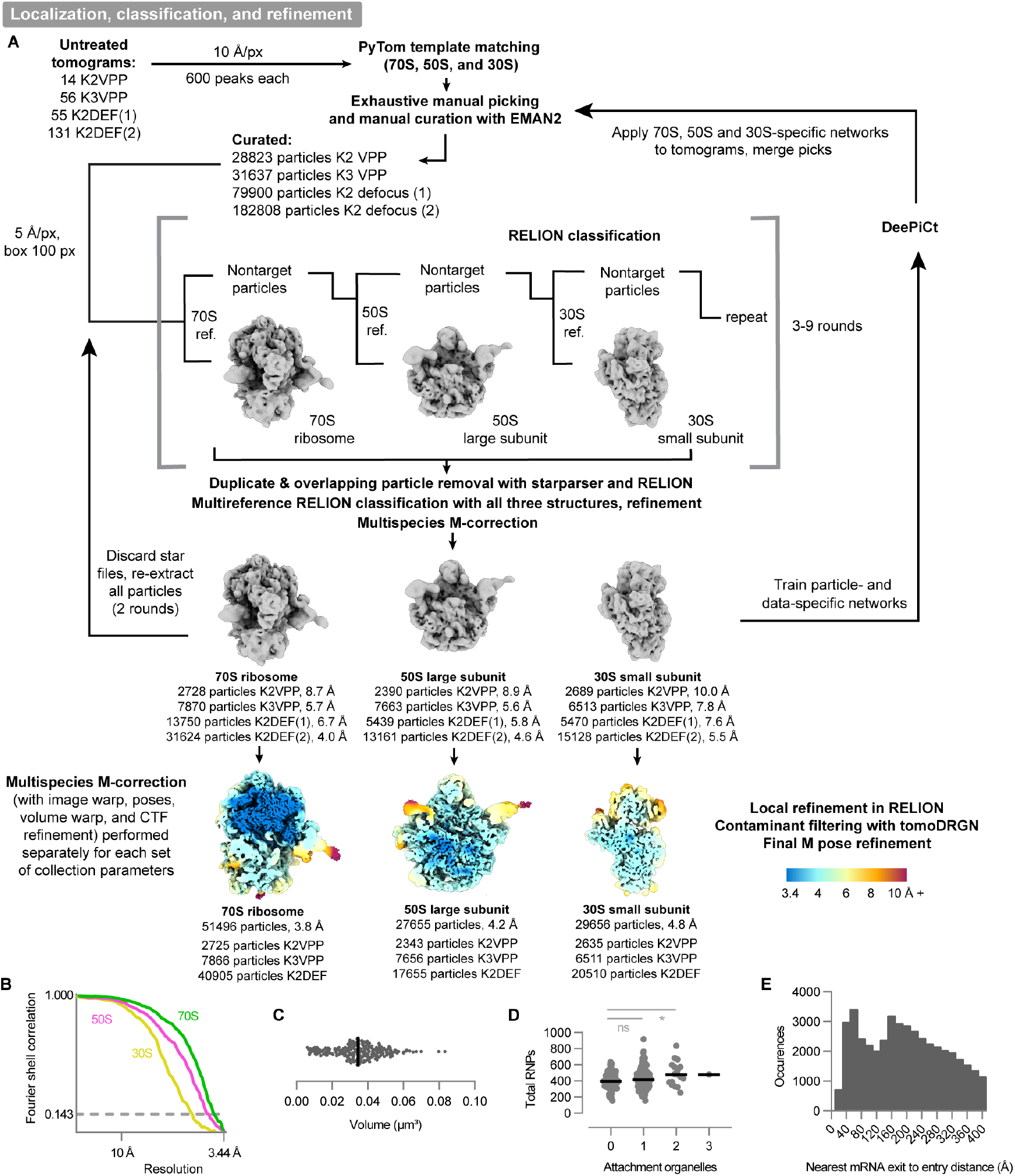
Particle picking, initial classification, and consensus refinement strategy. Related to Figure 1. **(A)** Particle picking and classification strategy as applied to native untreated *M. pneumoniae* cells, including local resolution maps for the consensus 70S (3.8 Å), 50S (4.2 Å), and 30S (4.8 Å) complexes. Local resolution is indicated in the colour bar (bottom right). Ref. indicates reference. **(B)** Fourier Shell Correlation (FSC) curves for the three consensus reconstructions. FSC threshold at 0.143 (dashed line) was used to determine the global resolution of the maps. **(C)** Volume distribution of analyzed *M. pneumoniae* cells in tomograms. Line indicates median: 0.035 μm^3^. **(D)** Plot of total RNP numbers in relation to the number of manually identified attachment organelles per cellular tomogram. **(E)** Histogram of distances of ribosome mRNA exit site to nearest ribosome mRNA entry site. Based on the histogram, a 70 Å cutoff was selected for polysome spatial analysis (Methods). Dots in plots represent individual cells and lines represent mean. Untreated (n=254) cells. By Student’s T test: *p < 0.05, **p < 0.01, ***p < 0.001, ns not significant.

**Figure S2:**
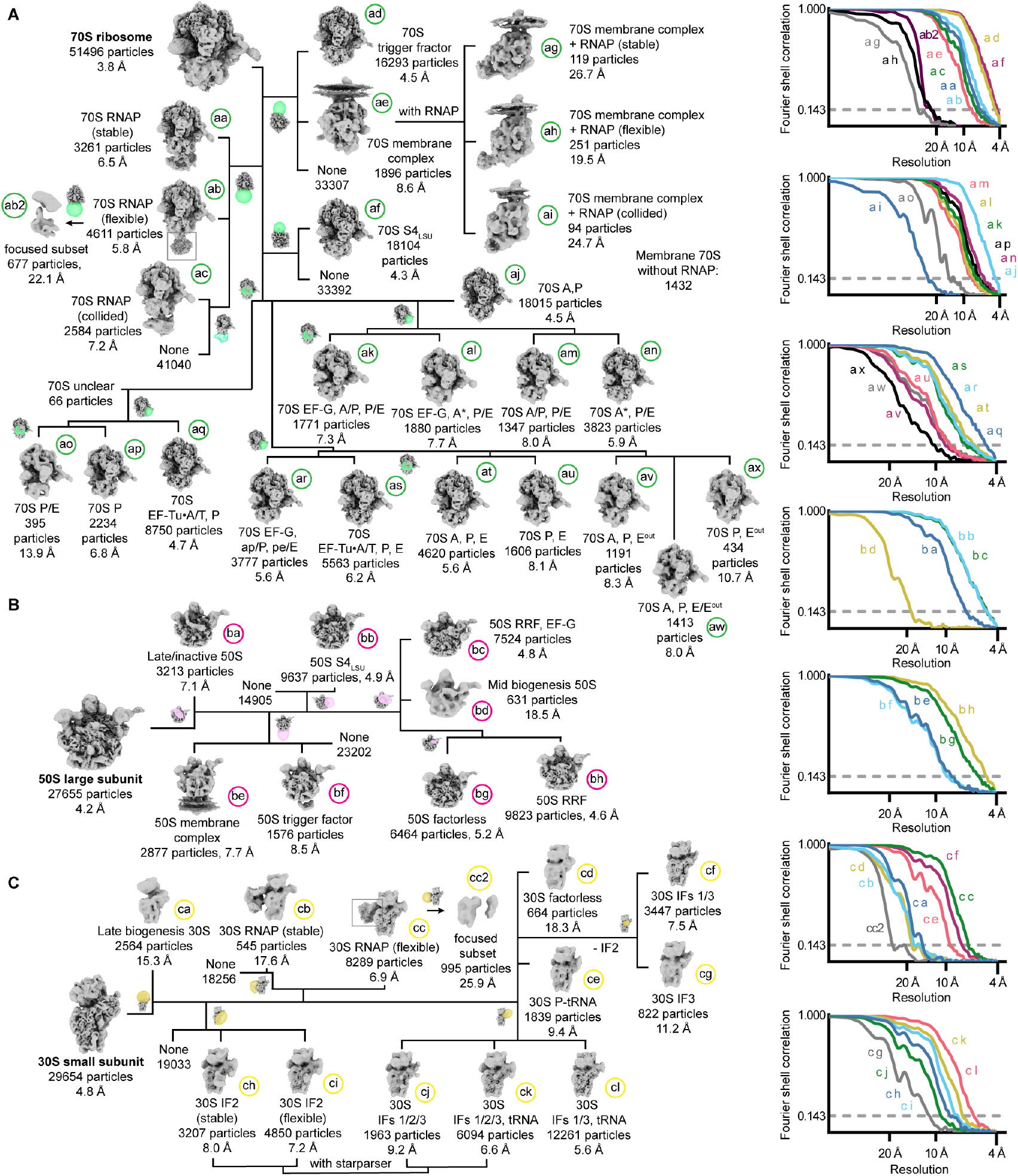
Classification of translation complexes (left) in native untreated *M. pneumoniae* cells, and corresponding FSC curves of final maps (right). Related to Figures 2-6. **(A)** Classification of 70S ribosomes. Local masks used in each classification are shown in green. Mask positioning was based on literature evidence, and multiple trials of positions and mask size. **(B)** Classification of 50S large subunits. Local masks used in each classification are shown in magenta. **(C)** Classification of 30S small subunits. Local masks used in each classification are shown in yellow. “None” indicates where complexes had no apparent extra density within the masked region.

**Figure S3:**
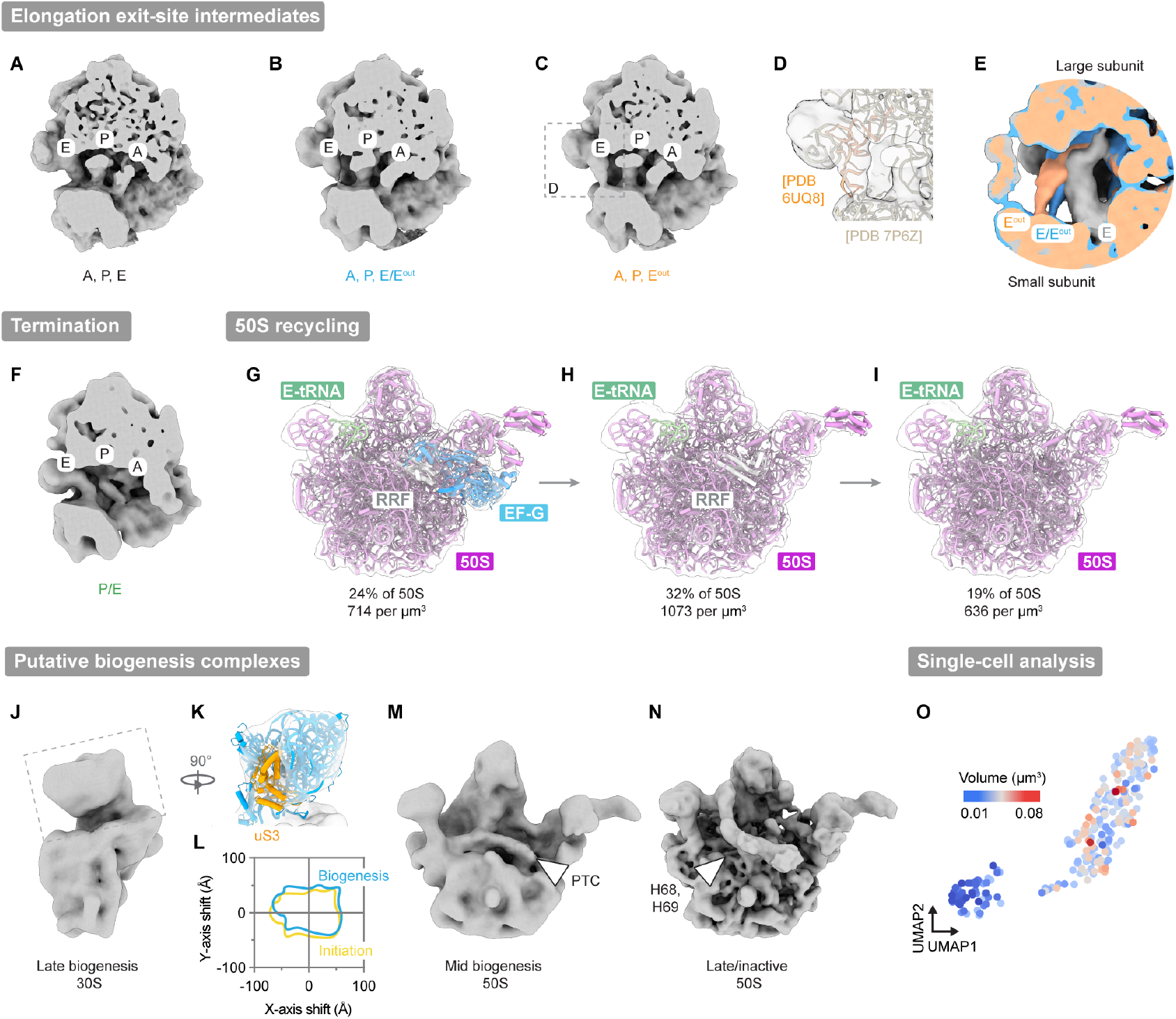
Additional maps in elongation, termination, and biogenesis. Related to Figures 2 and 3. **(A-C)** Maps of the 70S ribosome during elongation with different tRNA states: **(A)** Map of the 70S with A,P, and E tRNAs (5.6 Å). **(B)** Map of the 70S with A,P, and E/E^out^ tRNAs (8.0 Å). **(C)** Map of the 70S with A,P, and E^out^ tRNAs (8.3 Å). **(D)** Fit of the E^out^ tRNA from PDB 5UQ8^61^ to the experimental density (transparent) in the map shown in (C). **(E)** Comparison of E-site tRNA positions in **(A), (B)**, and **(C). (F)** Map of the 70S ribosome in the pre-recycling P/E state (13.9 Å). **(G-I)** Reconstructions depict the dissociation of factors from the 50S during ribosome recycling. Maps (transparent) are fitted with molecular models (cartoon) derived from combinations of experimental (PDB 7PAU, 7PAT)^123^ and predicted structures (Methods). **(G)** 50S with RRF and EF-G (4.8 Å). **(H)** 50S with RRF (4.6 Å). **(I)** Factorless 50S (5.2 Å). All 50S complexes were bound by a tRNA in the E-site. **(J-L)** Putative late 30S biogenesis complex: **(J)** Map of the complex (15.3 Å). **(K)** Fitting of 30S head region (blue, derived from PDB 7OOC)^45^ to the experimental density. uS3 (orange) is weak or absent in the map. **(L)** Comparison of fitted head position in consensus 30S complexes to the open head conformation in the putative biogenesis complex. **(M, N)** Maps of putative mid biogenesis and late/inactive 50S complexes: **(M)** Map of mid biogenesis 50S (18.5 Å) characterized by missing density in the peptidyl transferase centre (PTC, indicated by white arrowhead). **(N)** Map of late/inactive 50S (7.1 Å) characterized by incompletely folded rRNA helices 68 and 69 (indicated by white arrowhead). **(O)** Single-cell analysis visualized by UMAP (Methods). Projections is coloured by the volume of each cell (represented by individual dots).

**Figure S4:**
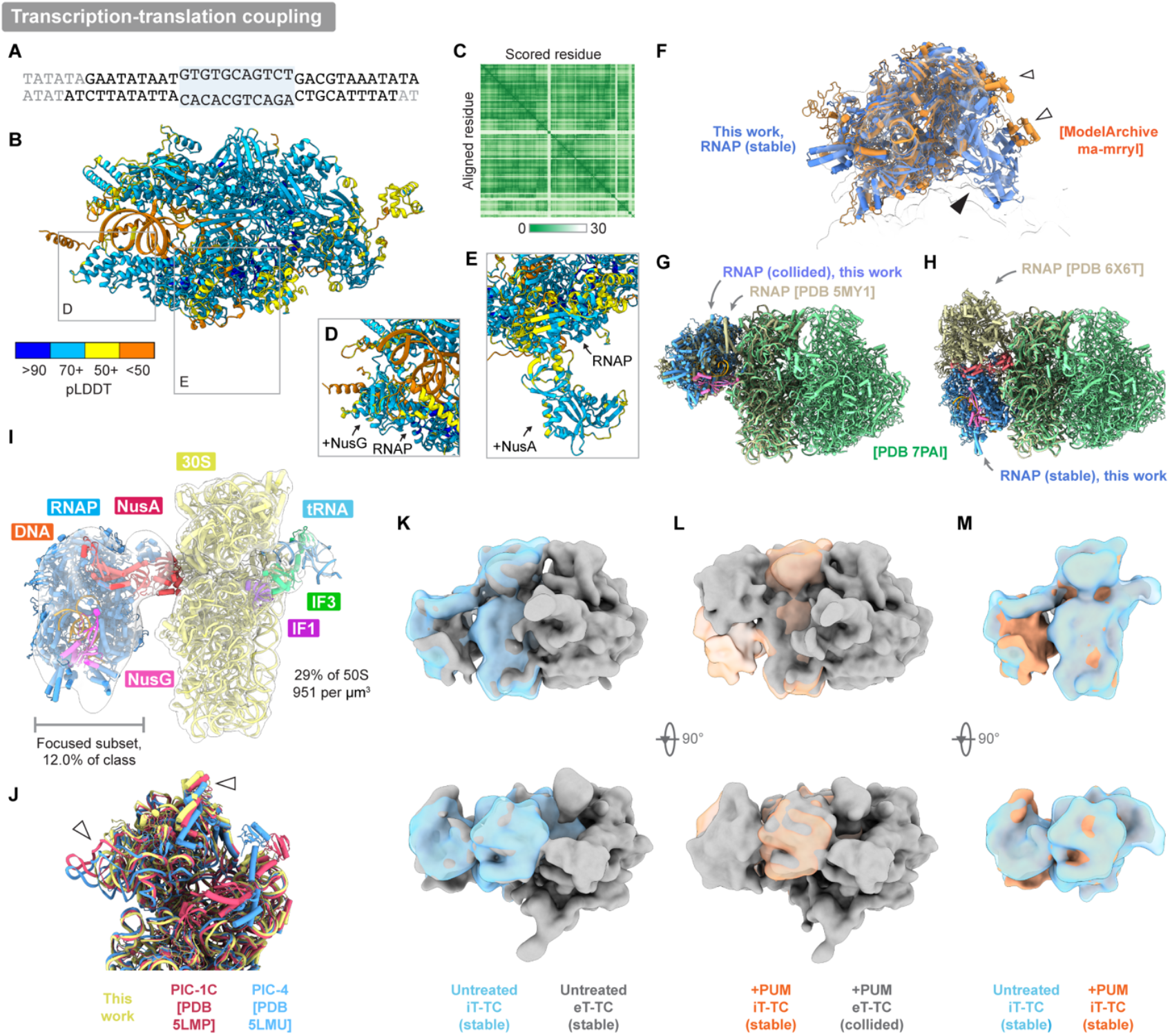
Molecular modeling of RNAP-associated translation complexes. Related to Figures 4 and 5. **(A)** DNA sequence (38 base pairs, adapted from ^144^) used for AlphaFold3 prediction of RNAP (Methods). Blue highlighting indicates the transcription bubble, bases in light gray are deleted from the final model. **(B)** AlphaFold3 prediction of the full *M. pneumoniae* RNAP (α, α, β, β’, δ) with DNA. Colour bar (bottom left) indicates corresponding predicted local distance difference test (pLDDT) score. **(C)** Plot of the predicted aligned error (PAE) of the AlphaFold3 RNAP model in (B). **(D, E)** AlphaFold3 prediction of RNAP with DNA and NusG **(D)**, or RNAP and NusA **(E)**, showing only the inset regions in (B). Scale indicates pLDDT score. **(F)** Comparison between the model from this work and a previous integrative model of the stable *M. pneumoniae* RNAP in native cells.^20^ Filled arrowhead indicates NusA as modelled here, and unfilled arrowheads indicate the relative repositioning of RNAP α C-terminal domains. Models are superimposed on the stable RNAP map from this work (transparent). **(G, H)** Comparisons between models from this work and previous studies. **(G)** Comparison between the collided eT-TC from this work and a model of a collided RNAP/70S complex from *E. coli*.^15^ **(H)** Comparison between the stable eT-TC from this work and a model of the *E. coli* RNAP bridged to the 70S via NusA and NusG.^17^ **(I)** Map and fitted model of the flexible iT-TC. A composite of maps, where RNAP was resolved with focused refinement of a subset of complexes and represents the highest likelihood conformation of RNAP in the flexible class. The focused subset RNAP map (25.9 Å) is shown alongside the consensus region (6.9 Å). **(J)** Comparison of 30S head position in the consensus 30S model here against various forms of initiating *T. thermophilus* 30S.^7^ **(K-M)** Comparisons between stable eT-TC and iT-TC maps low-pass filtered to 18 Å: **(K)** Untreated stable iT-TC and untreated stable eT-TC maps. **(L)** Untreated and PUM-treated stable iT-TC maps. **(M)** PUM-treated stable iT-TC and PUM-treated collided eT-TC maps.

**Figure S5:**
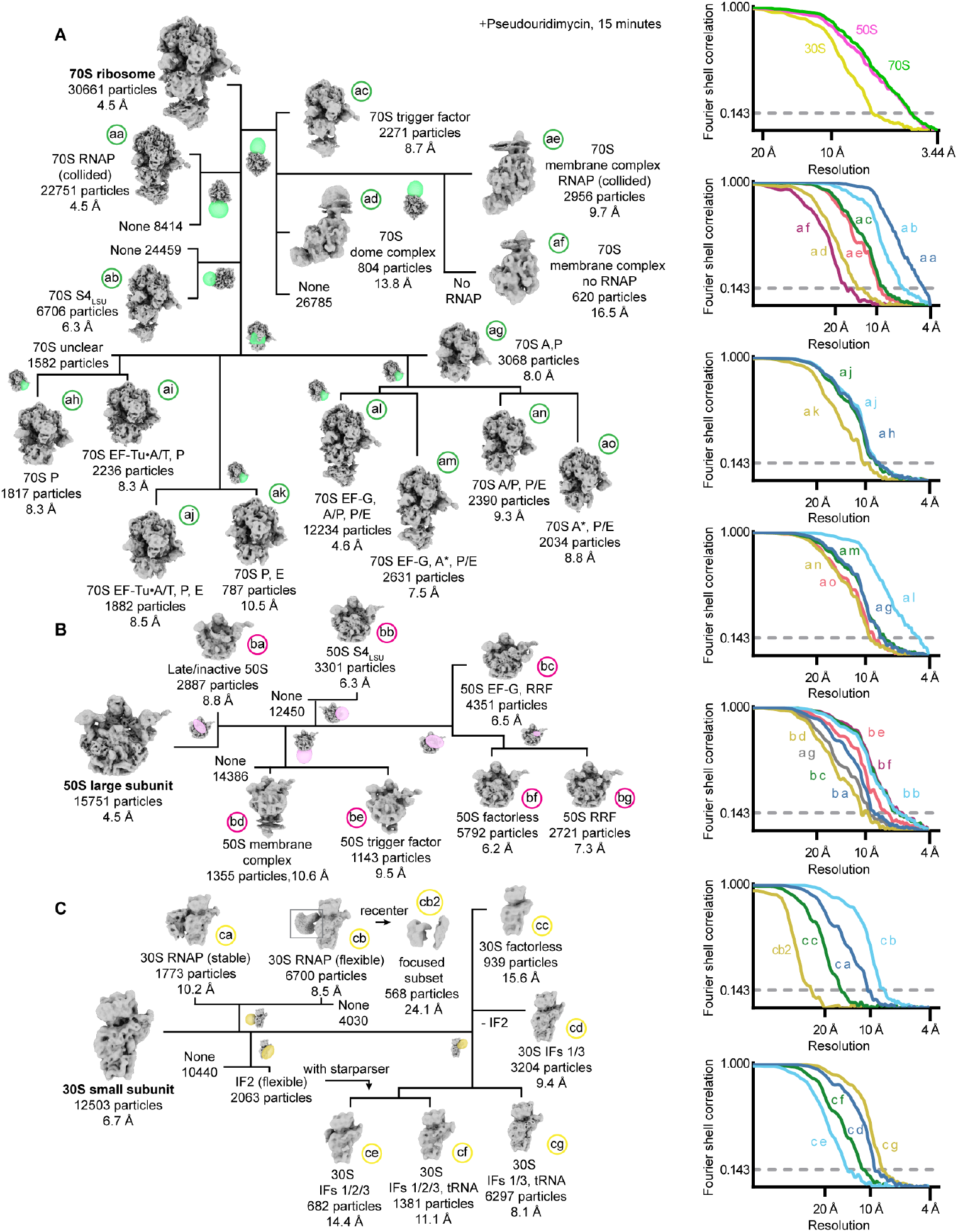
Classification of translation complexes in PUM-treated *M. pneumoniae* cells (left), and corresponding FSC curves of final maps (right). Related to Figures 5 and 6. **(A)** Classification of 70S ribosomes. Local masks used in each classification are shown in green. Mask positioning is described in Fig. S2A. **(B)** Classification of 50S large subunits. Local masks used in each classification are shown in magenta. **(C)** Classification of 30S small subunits. Local masks used in each classification are shown in yellow. “None” indicates where complexes had no apparent extra density within the masked region.

**Figure S6:**
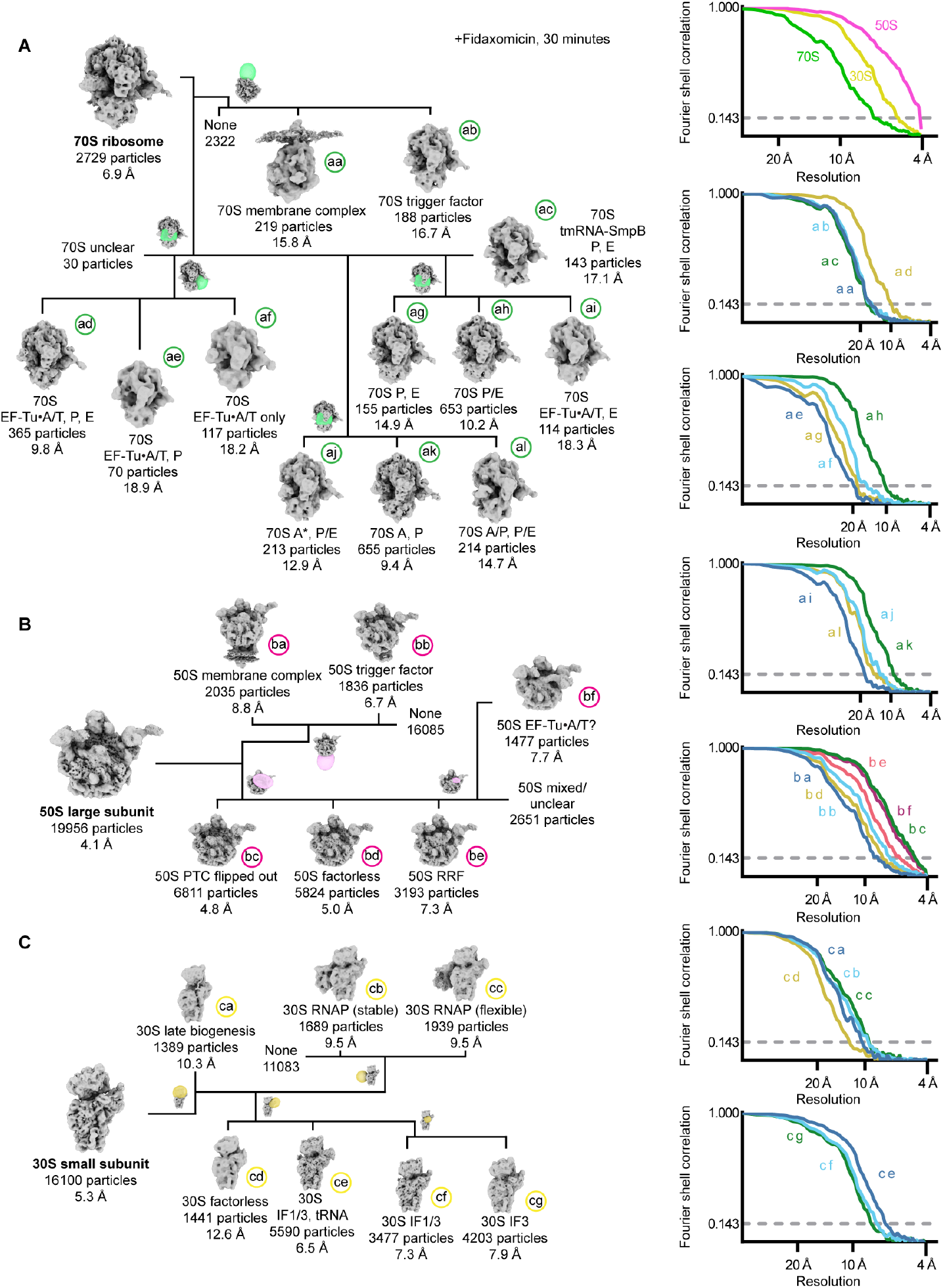
Classification of translation complexes in FID-treated *M. pneumoniae* cells (left), and corresponding FSC curves of final maps (right). Related to Figures 5 and 6. **(A)** Classification of 70S ribosomes. Local masks used in each classification are shown in green. Mask positioning is explained in Fig. S2A. **(B)** Classification of 50S large subunits. Local masks used in each classification are shown in magenta. **(C)** Classification of 30S small subunits. Local masks used in each classification are shown in yellow. “None” indicates where complexes had no apparent extra density within the masked region.

**Figure S7:**
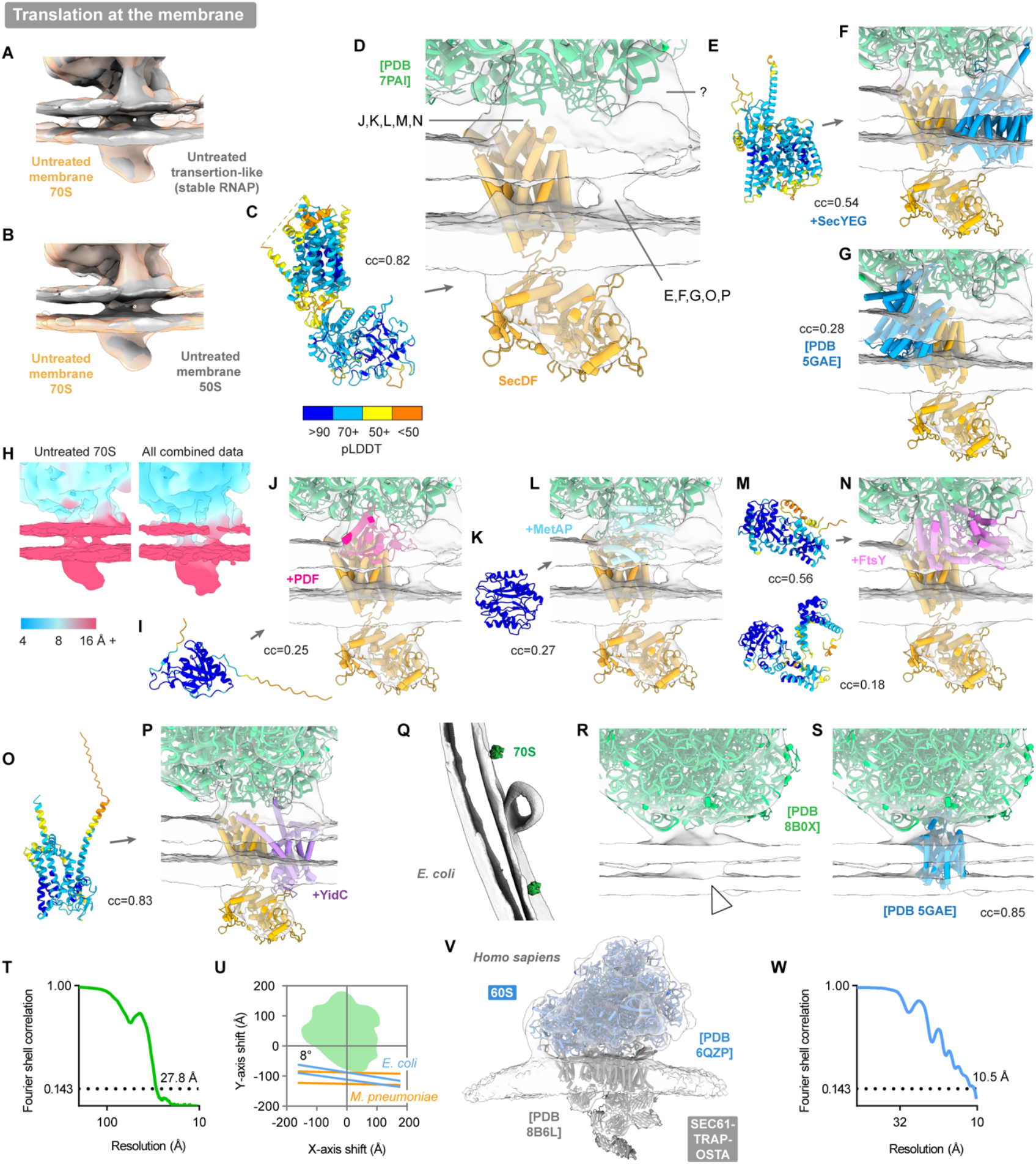
Molecular modeling of membrane-associated translation complexes. Related to Figure 6. **(A, B)** Untreated membrane-bound RNP maps low-pass filtered to 27 Å for comparison. Comparison between **(A)** membrane-bound 70S and transertion-like 70S+RNAP maps, or **(B)** membrane-bound 70S and membrane-bound 50S maps. **(C, D)** Rigid-body fit of an AlphaFold3 prediction of SecDF. **(C)** SecDF predicted structural model coloured by pLDDT score (mapped in colour bar, bottom), and fitted into the **(D)** membrane-bound 70S map together with a ribosome model (PDB 7PAI^45^). Unassigned densities are also indicated. cc indicates cross-correlation between the map and a simulation of the protein model density filtered to 20 Å resolution, which was performed in ChimeraX.^85^ **(E, F)** Rigid-body fit of an AlphaFold3 prediction of SecYEG coloured by pLDDT score **(E)** into unassigned density in the **(F)** membrane-bound 70S map. **(G)** Fit of an experimental model of the *E. coli* ribosome in complex with SecYEG (PDB 5GAE)^39^ to the membrane-bound 70S map (aligned based on ribosomal large subunit), with only SecYEG shown and focusing on the transmembrane region. Fit is inconsistent with the *M. pneumoniae* experimental density. **(H)** Transmembrane regions of untreated membrane 70S average and combined average of all membrane-associated complexes, coloured by local resolution (mapped in colour bar, bottom). **(I, J)** Rigid-body fit of an AlphaFold3 prediction of peptide deformylase (PDF) coloured by pLDDT score **(I)** into unassigned density in the **(J)** membrane-bound 70S map. **(K, L)** Rigid-body fit of an AlphaFold3 prediction of methionine aminopeptidase (MetAP) coloured by pLDDT score **(K)** into unassigned density in the **(L)** membrane-bound 70S map. **(M, N)** Rigid-body fits of AlphaFold3 predictions of the SRP receptor FtsY (top, fit shown) or SRP (bottom, fit not shown) coloured by pLDDT score **(M)** into unassigned density in the **(N)** membrane-bound 70S map. **(O, P)** Rigid-body fit of an AlphaFold3 prediction of YidC coloured by pLDDT score **(O)** into unassigned density in the **(P)** membrane-bound 70S map. **(Q-T)** Analysis of in-cell *E. coli* 70S ribosomes bound to the membrane. **(Q)** Membrane segmentation and localization of two membrane-bound 70S ribosomes. **(R)** Map of the membrane bound 70S in *E. coli* (27.8 Å), focusing on the transmembrane region, fit with PDB 8B0X.^121^ White arrowhead indicates transmembrane density. **(S)** Aligned fit (based on ribosomal large subunit) of the *E. coli* ribosome in complex with SecYEG (PDB 5GAE)^39^ to the map. **(T)** FSC curve corresponding to the membrane-bound *E. coli* 70S reconstructed at 5 Å/px. **(U)** Comparison of the ribosome-membrane attachment angle for *M. pneumoniae* and *E. coli*. Membrane angle differs by 8°. **(V, W)** Structural analysis of the human ER-bound large subunit. **(V)** Map of a membrane-bound human 60S large subunit with SEC61-TRAP-OSTA translocon (10.5 Å) derived from re-analysis of EMPIAR-11751 (Methods)^50^ and fitted with PDBs 6QZP^122^ and 6B6L.^50^ **(W)** Corresponding FSC curve for map data, reconstructed at 5 Å/px.

**Figure S8:**
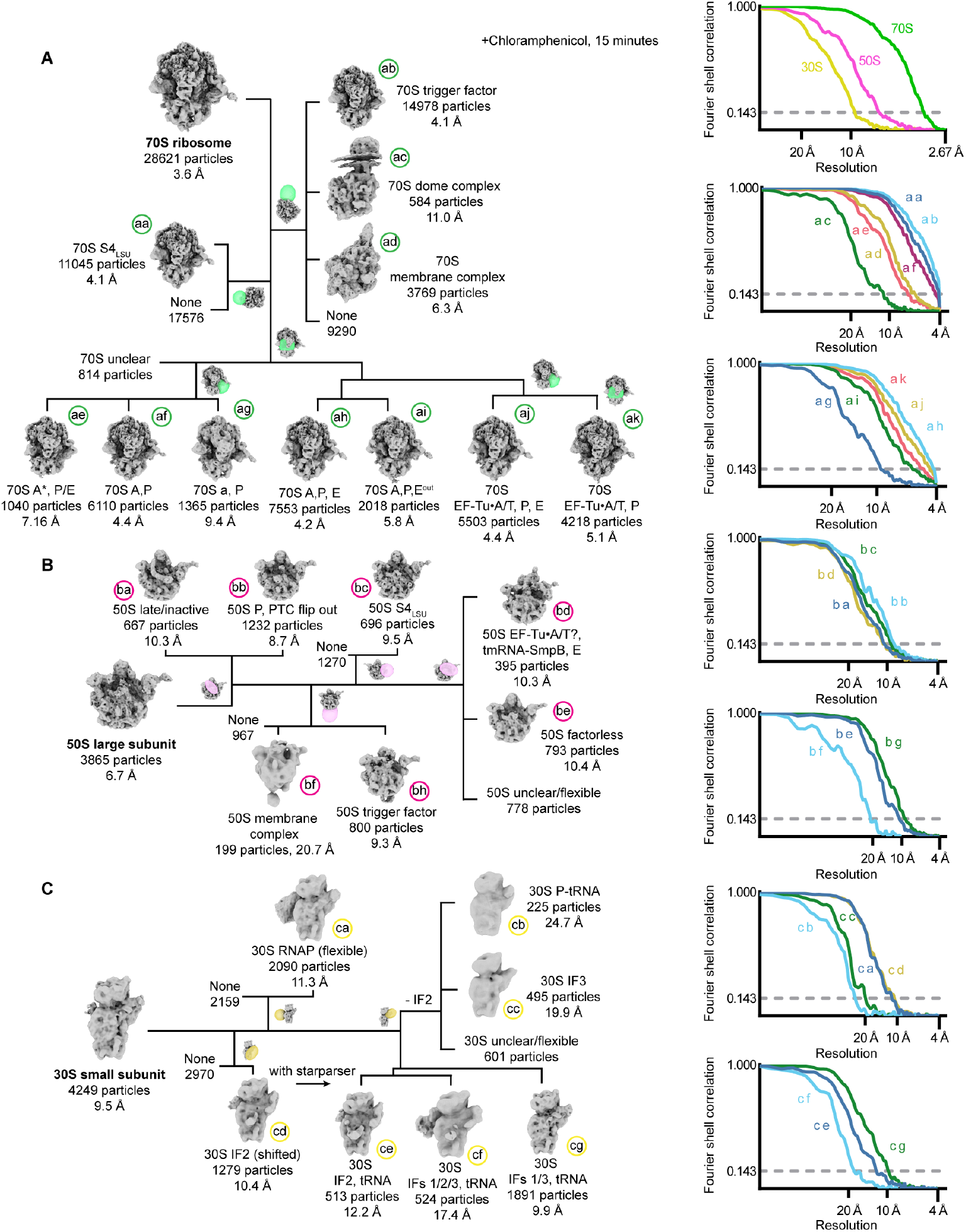
Classification of translation complexes in Cm-treated *M. pneumoniae* cells (left), and corresponding FSC curves of final maps (right). Related to Figure 6. **(A)** Classification of 70S ribosomes. Local masks used in each classification are shown in green. Mask positioning is explained in Fig. S2A. **(B)** Classification of 50S large subunits. Local masks used in each classification are shown in magenta. **(C)** Classification of 30S small subunits. Local masks used in each classification are shown in yellow. Of note: IF2 and the initiator tRNA are capable of binding the 30S without IF1 in this condition, and IF2’s position is shifted, possibly due to a relative increase in concentrations of these in comparison to 30S complexes. “None” indicates where complexes had no apparent extra density within the masked region.

**Table S1:** *M. pneumoniae* translation complexes detailed across conditions. Related to figures 1-6. # represents total number of complexes, % is the fraction out of all RNPs, and Å the global resolution of the map at FSC threshold 0.143. * indicates where a structural model was generated and deposited. – indicates where a final refined model was not generated, but where we report the number of complexes (*e*.*g*. unclear identity, or obtained by taking the overlap between two particle sets).

| Condition | Untreated |  |  | PUM |  |  | FID |  |  | Cm |  |  |
| --- | --- | --- | --- | --- | --- | --- | --- | --- | --- | --- | --- | --- |
| Magnification | 81000x and 53000x |  |  | 53000x |  |  | 53000x |  |  | 64000x |  |  |
| Voltage (kV) | 300 |  |  | 300 |  |  | 300 |  |  | 300 |  |  |
| Electron exposure (e<sup>-</sup>/Å<sup>2</sup>) | ~120 |  |  | ~120 |  |  | ~120 |  |  | ~120 |  |  |
| Defocus range (µm) | -2 to -5 |  |  | -2 to -4 |  |  | -2 to -5 |  |  | -2 to -4 |  |  |
| Pixel size (Å/pixel) | 1.71 and 1.631 |  |  | 1.631 |  |  | 1.631 |  |  | 1.329 |  |  |
| Tomogram/cell number | 254 |  |  | 124 |  |  | 100 |  |  | 121 |  |  |
| Total RNPs | 108805 |  |  | 58915 |  |  | 38785 |  |  | 36735 |  |  |
| Complex | # | % | Å | # | % | Å | # | % | Å | # | % | Å |
| 30S P-tRNA* | 1839 | 1.7 | 9.4 |  |  |  |  |  |  | 225 | 0.6 | 24.7 |
| 30S IF3* | 822 | 0.8 | 11.2 |  |  |  | 4203 | 10.8 | 7.9 | 495 | 1.3 | 19.9 |
| 30S IF1/3* | 3447 | 3.2 | 7.5 | 3204 | 5.4 | 9.4 | 3477 | 8.9 | 7.3 |  |  |  |
| 30S empty/factorless* | 664 | 0.6 | 18.3 | 939 | 1.6 | 15.6 | 1441 | 3.7 | 12.6 |  |  |  |
| 30S IF1/2/3* | 1963 | 1.8 | 9.2 | 682 | 1.2 | 14.4 |  |  |  |  |  |  |
| 30S IF1/3, tRNA* | 12261 | 11.3 | 5.6 | 6296 | 10.7 | 8.1 | 5590 | 14.4 | 6.5 | 1891 | 5.1 | 9.9 |
| 30S IF 1/2/3, tRNA* | 6094 | 5.6 | 6.6 | 1381 | 2.3 | 11.1 |  |  |  | 524 | 1.4 | 17.4 |
| 30S iT-TC flexible* | 8289 | 7.6 | 6.9 | 6700 | 11.4 | 8.5 | 1939 | 5.0 | 9.5 | 2090 | 5.7 | 11.3 |
| 30S iT-TC stable** | 545 | 0.5 | 17.6 | 1773 | 3.0 | 10.2 | 1689 | 4.3 | 9.5 |  |  |  |
| 30S flexible IF2 (all) | 4850 | 4.5 | 7.2 | 2063 | 3.5 | - |  |  |  |  |  |  |
| 30S stable IF2 (all) | 3207 | 2.9 | 8.0 |  |  |  |  |  |  |  |  |  |
| 30S IF2 (shifted), all |  |  |  |  |  |  |  |  |  | 1279 | 3.5 | 10.4 |
| 30S IF2 (shifted), P-tRNA |  |  |  |  |  |  |  |  |  | 513 | 1.4 | 12.2 |
| Late biogenesis 30S | 2564 | 2.4 | 15.3 |  |  |  | 1389 | 3.6 | 10.3 |  |  |  |
| 30S unclear |  |  |  |  |  |  |  |  |  | 601 | 1.6 | - |
| 50S RRF, EF-G* | 7524 | 6.9 | 7.1 | 4351 | 7.4 | 6.5 |  |  |  |  |  |  |
| 50S RRF | 9823 | 9.0 | 4.6 | 2721 | 4.6 | 7.3 | 3193 | 8.2 | 7.3 |  |  |  |
| 50S empty/factorless | 6464 | 5.9 | 5.2 | 5792 | 9.8 | 6.2 | 5824 | 15.0 | 5.0 | 793 | 2.2 | 10.4 |
| 50S S4<sub>LSU</sub> | 9637 | 8.9 | 4.9 | 3301 | 5.6 | 6.3 |  |  |  | 696 | 1.9 | 9.5 |
| 50S trigger factor | 1576 | 1.4 | 8.5 | 1143 | 1.9 | 9.5 | 1863 | 4.8 | 6.7 | 800 | 2.2 | 9.3 |
| Membrane-bound 50S* | 2877 | 2.6 | 7.7 | 1355 | 2.3 | 10.6 | 2035 | 5.2 | 8.8 | 199 | 0.5 | 20.7 |
| Mid biogenesis 50S | 631 | 0.6 | 18.5 |  |  |  |  |  |  |  |  |  |
| Late/inactive 50S | 3213 | 3.0 | 7.1 | 2887 | 4.9 | 8.8 |  |  |  | 667 | 1.8 | 10.3 |
| 50S PTC flipped out |  |  |  |  |  |  | 6811 | 17.6 | 4.8 |  |  |  |
| 50S PTC flipped out, P |  |  |  |  |  |  |  |  |  | 1232 | 3.4 | 8.7 |
| 50S EF-Tu A/T? |  |  |  |  |  |  | 1477 | 3.8 | 7.7 |  |  |  |
| 50S EF-Tu A/T?, tmRNA-SmpB,P,E |  |  |  |  |  |  |  |  |  | 395 | 1.1 | 10.3 |
| 50S unclear |  |  |  |  |  |  |  |  |  | 778 | 2.1 | - |
| 70S P | 2234 | 2.1 | 6.8 | 1817 | 3.1 | 8.3 |  |  |  |  |  |  |
| 70S P,E | 1606 | 1.5 | 8.1 | 787 | 1.3 | 10.5 | 155 | 0.4 | 14.9 |  |  |  |
| 70S EF-Tu A/T,P,E | 5563 | 5.1 | 6.2 | 1882 | 3.2 | 8.5 | 365 | 0.9 | 9.8 | 5503 | 15.0 | 4.4 |
| 70S EF-Tu A/T |  |  |  |  |  |  | 117 | 0.3 | 18.2 |  |  |  |
| 70S A,P,E | 4620 | 4.2 | 5.6 |  |  |  |  |  |  | 7553 | 20.6 | 4.2 |
| 70S A,P,E/E<sup>out</sup> | 1413 | 1.3 | 8.0 |  |  |  |  |  |  |  |  |  |
| 70S A,P,E<sup>out</sup> | 1191 | 1.1 | 8.3 |  |  |  |  |  |  | 2018 | 5.5 | 5.8 |
| 70S P,E<sup>out</sup> | 434 | 0.4 | 10.7 |  |  |  |  |  |  |  |  |  |
| **70S EF-Tu A/T, P** | 8750 | 8.0 | 4.7 | 2236 | 3.8 | 8.3 | 70 | 0.2 | 18.9 | 4218 | 11.5 | 5.1 |
| **70S EF-Tu A/T, E** |  |  |  |  |  |  | 114 | 0.3 | 18.3 |  |  |  |
| **70S A,P** | 12626 | 11.6 | 4.5 | 3068 | 5.2 | 8.0 | 655 | 1.7 | 9.4 | 6110 | 16.6 | 4.4 |
| **70S A\*, P/E** | 3823 | 3.5 | 7.7 | 2034 | 3.5 | 8.8 | 213 | 0.6 | 12.9 | 1040 | 2.8 | 7.2 |
| **70S A/P, P/E** | 1347 | 1.2 | 8.0 | 2390 | 4.1 | 9.3 | 214 | 0.6 | 14.7 |  |  |  |
| **70S EF-G A\*, P/E** | 1880 | 1.7 | 7.7 | 2631 | 4.5 | 7.5 |  |  |  |  |  |  |
| **70S EF-G A/P, P/E** | 1771 | 1.6 | 7.3 | 12234 | 20.8 | 4.6 |  |  |  |  |  |  |
| **70S EF-G ap/P pe/E** | 3777 | 3.5 | 5.6 |  |  |  |  |  |  |  |  |  |
| **70S P/E** | 395 | 0.4 | 13.9 |  |  |  | 653 | 1.7 | 10.2 |  |  |  |
| **70S S4<sub>LSU</sub>** | 18104 | 16.6 | 4.3 | 6706 | 11.4 | 6.3 |  |  |  | 11045 | 30.1 | 4.1 |
| **Membrane-bound 70S\***<br>( ) = not including +RNAP | 1896<br>(1432) | 1.7<br>(1.3) | 8.6<br>- | 3576<br>(620) | 6.1<br>(1.1) | -<br>(17) | -<br>(219) | -<br>(0.6) | -<br>(16) | -<br>(3769) | -<br>(10) | -<br>(6.3) |
| **70S trigger factor** | 16293 | 15.0 | 4.5 | 2271 | 3.9 | 8.7 | 188 | 0.5 | 16.7 | 14978 | 40.8 | 4.1 |
| **70S eT-TC flexible\*** | 4611 | 4.2 | 5.8 |  |  |  |  |  |  |  |  |  |
| **70S eT-TC stable\*** | 3261 | 3.0 | 6.5 |  |  |  |  |  |  |  |  |  |
| **70S eT-TC collided\*** | 2584 | 2.4 | 7.2 | 22751 | 38.6 | 4.5 |  |  |  |  |  |  |
| **Membrane-bound 70S flexible RNAP** | 251 | 0.2 | 19.5 |  |  |  |  |  |  |  |  |  |
| **Membrane-bound 70S stable RNAP\*** | 119 | 0.1 | 26.7 |  |  |  |  |  |  |  |  |  |
| **Membrane-bound 70S collided RNAP** | 94 | 0.09 | 24.7 | 2956 | 5.0 | 9.7 |  |  |  |  |  |  |
| **70S dome complex** |  |  |  | 804 | 1.4 | 13.8 |  |  |  | 584 | 1.6 | 11.0 |
| **70S tmRNA-SmpB, P,E** |  |  |  |  |  |  | 143 | 0.4 | 17.1 |  |  |  |
| **70S a,P** |  |  |  |  |  |  |  |  |  | 1365 | 3.7 | 9.4 |
| **70S tRNA unclear** | 66 | 0.1 | - | 1582 | 2.7 | - | 30 | 0.1 | - | 814 | 2.2 | - |

